# OPUS-ET: Resolving Compositional and Conformational Heterogeneities of Biomolecules in Cryo-Electron Tomography

**DOI:** 10.1101/2025.11.21.688990

**Authors:** Zhenwei Luo, Xiangru Chen, Qinghua Wang, Jianpeng Ma

## Abstract

Structural heterogeneity in biomolecules, arising from both compositional and conformational variability, limits resolution and interpretability of cryo-electron tomography (cryo-ET). Here, we present OPUS-ET, a deep learning framework that resolves multiscale heterogeneity throughout the cryo-ET workflow. OPUS-ET combines a composition decoder that captures compositional differences with a conformation decoder that models large-scale motions, thereby providing a hierarchical representation of structural heterogeneity. Starting from noisy template-matching candidates with templates of varying similarity or quality, OPUS-ET efficiently enriches target particle populations and delivers sub-nanometer *in situ* reconstructions in a single round. It leads to improved resolutions by up to 4.5 Å over expert annotations or existing deep-learning approaches in four benchmark systems, and reveals continuous conformational landscapes capturing F₀–F₁ flexible coupling in mitochondrial ATP synthase and tRNA-translocation intermediates in eukaryotic and bacterial ribosomes. Together, these results establish OPUS-ET as a powerful computational tool for linking particle purification, high-resolution reconstruction, and analysis of structural heterogeneity in cryo-ET, with demonstrated robustness to template quality, initial pose noise, and clustering parameters.

## Introduction

Recent advances in single-tilt axis holders^1^, cryo-focused ion beam (cryo-FIB) instruments^2,3^, and image processing algorithms^4–8^ have significantly propelled cryo-electron tomography (cryo-ET) toward sub-nanometer resolution^4^. Unlike cryo-electron microscopy (cryo-EM), cryo-ET preserves macromolecules within their native cellular environment^9,10^. It captures three-dimensional (3D) snapshots of “molecular sociology”^11^ and resolves intermediate states *in situ* that are inaccessible to conventional methods^12^. Yet, resolving the full complexity of in-cell macromolecular structures remains hampered by multi-scale structural heterogeneity.

Structural heterogeneity in cryo-ET arises from both compositional and conformational variabilities^13,14^. Low-dose imaging, missing-wedge artifacts, defocus variations, and crowded cellular environments collectively produce low signal-to-noise ratios (SNRs) ^9–11,15^. As a result, template-based or template-free particle-picking approaches inevitably include false positives, and even extensively classified subtomogram sets retain substantial heterogeneity^16–20^. These heterogeneities, although often biologically meaningful, limit achievable resolution and complicate functional interpretation^10,11,21^. An integrated framework capable of resolving both compositional and conformational variabilities across the entire cryo-ET workflow is therefore essential for high-resolution, mechanistically interpretable reconstructions.

Existing software such as pyTOM, RELION, and nextPYP separate compositional variants through hierarchical 3D classification but are restricted to uncovering a few discrete states^5,7,9,22,23^. Various methods have been designed to model continuous conformational variations using different structured descriptions of molecular motions, including multi-body refinement (RELION^24^), neural-network-based representations (cryoSPARC 3D Flex^25^,

DynaMight^26^, Gaussian mixture model^27^), normal-mode basis (WarpCraft^28^) and spherical-harmonic or Zernike-based representations (Zernike3D^29,30^). Recent deep-learning methods (CryoDRGN^31^ and OPUS-DSD^32^) introduced neural-network–based 3D volume representations and inspired cryo-ET adaptations like tomoDRGN^33^ and CryoDRGN-ET^34^. However, their volume-based parameterizations remain inefficient for large-scale dynamics and entangle compositional and conformational variabilities within a single latent space.

Here, we introduce OPUS-ET, a deep learning framework that simultaneously resolves compositional and conformational heterogeneity throughout the cryo-ET workflow. OPUS-ET encodes each subtomogram into two latent spaces: a **composition latent space** decoded by a 3D convolutional neural network (CNN) to capture what is present, and a **conformation latent space** decoded by a physics-informed multi-layer perceptron (MLP) to model inter-subunit dynamics. This separation enables hierarchical modeling of compositional variability and inter-subunit dynamics within a single framework. In a series of benchmark systems, OPUS-ET produces high-resolution reconstructions that outperform expert annotation and existing deep-learning approaches by up to 4.5 Å after M-refinement. The reconstructions reveal continuous conformational landscapes, examples include the F₀–F₁ flexible coupling in ATP synthase and tRNA translocation in ribosomes. By automating heterogeneity analysis and linking particle purification with analysis of structural variation *in situ*, OPUS-ET efficiently bridges high-resolution studies of molecular structures and functions in cells.

## Results

### Neural architecture of OPUS-ET

Template matching and subtomogram averaging are two essential steps in cryo-ET structural determination. Template matching identifies the position and orientation of macromolecules within a tomogram, whereas subtomogram averaging refines the particles relative to a consensus reference. Both steps produce particle sets with estimated poses, but inevitably retain substantial structural heterogeneity, from compositional difference to conformational variations, which OPUS-ET is designed to resolve.

OPUS-ET accepts inputs from template-matching or subtomogram averaging results. Each subtomogram is aligned to the canonical orientation of its reference and then passed through an encoder that estimates the distributions of two latent codes (**Fig.1a**): a composition latent code 𝑧 and a conformation latent code 𝑧*_d_*. For each latent code, the encoder predicts the mean 𝑧 ∈ ℝ*^n^* and standard variation 𝜎 ∈ ℝ*^n^* ^35^. Latent samples are drawn from these distributions and supplied to their respective decoders.

**Figure 1.**
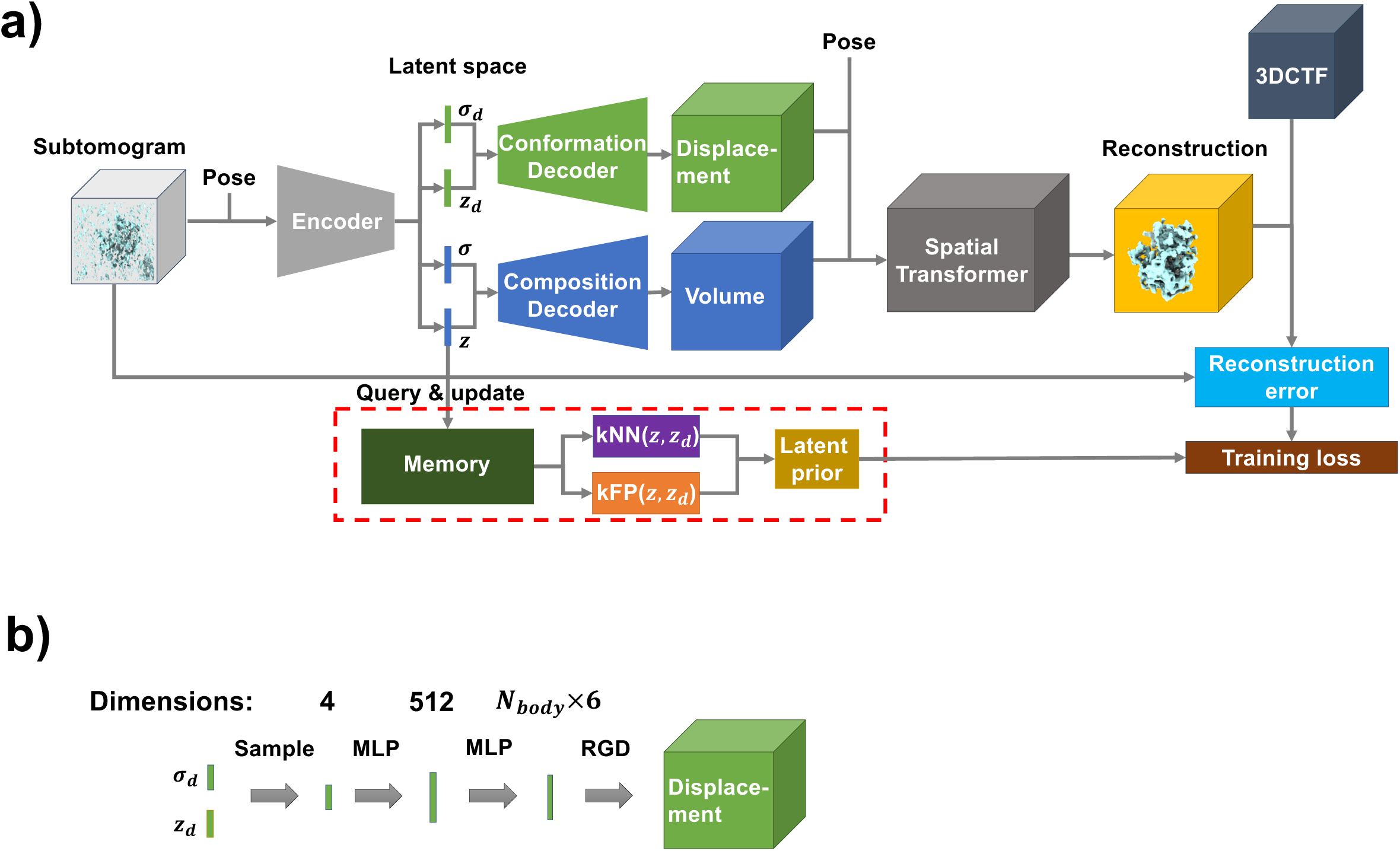
OPUS-ET architecture. **a)**. OPUS-ET workflow linking input subtomograms to reconstructed 3D maps through disentangled composition and conformation latent spaces. Pose denotes each subtomogram’s orientation relative to the consensus reference. 3DCTF denotes per-particle contrast-transfer function. 𝑧 and 𝜎 are the mean and standard deviation vectors of composition latent code, respectively. 𝑧_!_ and 𝜎_!_ are the mean and standard deviation vectors of conformation latent code, respectively. kNN(𝑧, 𝑧_!_) and kFP(𝑧, 𝑧_!_) refer to the K nearest neighbor and K farthest points of the concatenation of 𝑧 and 𝑧_!_, respectively. Auxiliary regularization connections are omitted for clarity. **b)**. Conformation decoder translating latent vectors 𝑧_!_into 3D deformation fields through multi-layer perceptrons (MLPs). Rigid-body dynamics (RGD) module converts predicted rotations and translations into the 3D deformation field. All MLPs except the last one use LeakyReLU (non-linearity = 0.2).

The composition latent code is processed by a 3D convolutional decoder that reconstructs a voxelized 3D volume 𝑉(𝒙) to represent the structural content of the subtomogram (**Fig.1a**). In parallel, the conformation latent code is decoded by an MLP into a set of rigid-body transformations, one for each predefined subunit. These transformations are smoothly blended into a deformation field describing inter-subunit motion. OPUS-ET uses this field in an inverse-mapping manner: for each output voxel coordinate 𝒙 in the deformed volume, the model predicts the corresponding source coordinate in the reference volume 𝑉(𝒙), and the output intensity is obtained by trilinear interpolation at that source location through a spatial transformer^36^. In this way, the deformation field and the reference volume 𝑉(𝒙) are combined to generate the final 3D reconstruction at the estimated pose (**Fig.1a**). This structure is then converted into a synthetic subtomogram by incorporating the corresponding 3D-contrast transfer function (3DCTF) using the differentiable imaging model described in **Methods**.

OPUS-ET is trained end-to-end by minimizing the squared reconstruction error between each experimental and reconstructed subtomogram. During training, a query-and-update module is used to iteratively refine the latent representation associated with each subtomogram across randomly-shifted mini-batches through momentum averaging (**Fig.1a**). Both latent spaces are regularized with the latent prior from OPUS-DSD^32^ and a Kullback-Leibler (KL) divergence term towards a standard Gaussian^35,37^ (**Fig.1a**), encouraging smooth and well-structured latent manifolds. This design enables OPUS-ET to separate conformational and compositional heterogeneities directly from noisy tomograms.

The neural network design for each pathway is explained as follows. The encoder and composition decoder follow the OPUS-DSD 3D CNN design^32^, while the conformation decoder implements a nonlinear MLP (**Fig.1b**). The conformation latent code is mapped to a 512-dimensional representation via LeakyReLU layers and the projected to an 𝑁_body_ × 6-dimensional vector, where 𝑁_body_denotes the number of predefined rigid bodies. Each 𝑁_body_ × 6-dimensional vector encodes rotation (quaternion) and translation of one subunit around its center of mass. The resulting deformation field is smoothly blended from these transformations (see **Methods**).

### Purification and high-resolution reconstruction of *Chlamydomonas reinhardtii* mitochondrial ATP synthase

ATP synthase is a mitochondrial protein complex that synthesizes ATP using the proton-motive force across the inner membrane. Previous *in situ* cryo-ET studies resolved the *Polytomella* dimer at 6.8 Å^23^ and sub-nanometer reconstructions of *Chlamydomonas (C.) reinhardtii* peripheral stalk and F₁-head, but lacked a full-dimer reconstruction^38^. These analyses required repeated, expert-driven 3D classifications that are labor-intensive and time-consuming.

Using pyTOM^7^ template matching with the 15Å low-pass filtered *Polytomella* ATP synthase map^23^ (EMDB code EMD-50000) as reference, we extracted 783,000 subtomograms from 261 *C. reinhardtii* mitochondrial tomograms and performed two filtering rounds. In the first round, subtomograms from 62 random tomograms were used to train an initial OPUS-ET model to remove false positives such as membranes. By UMAP visualization of learned composition latent space, among 20 latent-space classes, Class 13 (10,722 subtomograms) contained complete dimers, Class 15 (43,173 subtomograms) exhibited weaker densities, and remaining classes were dominated by membranes and noise (**Ext. Data Fig.1a**). The remaining 199 tomograms were similarly processed, and classes similar to Classes 13 and 15 in **Ext. Data Fig.1a** were combined, yielding 220,291 subtomograms, accounting for 28% of all template-matching picks (220,291/783,000).

In the second round, OPUS-ET further separated ATP synthase from background and revealed higher-order oligomers (**Ext. Data Fig.1b**). Class 2 represented dimers with faint neighboring densities, Classes 1 and 4 corresponded to tetramers, and Class 3 captured hexamers interlocked *via* ATP synthase-associated protein (ASA) subunits (**Fig.2a**), consistent with *Polytomella in situ* structures^23^. Refinement of Classes 1∼4 (a total of 34,367 subtomograms) in M^4^ focusing on the central dimer with C_2_ symmetry produced an 8.6 Å average (**Fig.2b; Ext. Data Fig.2a**). Including 15,354 additional subtomograms from the top dimers in Classes 3 and 4 (totally 49,721 particles, termed as OPUS-ET Expanded) slightly improved the nominal resolution to 8.5 Å, but substantially enhanced local densities (**Fig.2c**). Compared with the 6.8 Å *Polytomella* dimer^23^ (**Fig.2d**), our 8.5-Å *C. reinhardtii* map showed similar helical features but lacked the ASA “bridge” of the peripheral stalk, a key interspecies difference^38^.

To benchmark OPUS-ET against human expert annotation, we re-analyzed the *C. reinhardtii* dataset from Ron *et al*.^38^ (150,864 subtomograms). Refinement of this expert-annotated set yielded an 11.9 Å map without helical features (**Fig.2e**; **Ext. Data Fig.2a**). Applying OPUS-ET to this dataset identified Classes 3∼13 (totally 56,194 subtomograms, termed as Expert annotations + OPUS-ET) as ATP synthase oligomers (**Ext. Data Fig.1c**). Refinement reached 9.1 Å with clear helical features (**Fig.2f**; **Ext. Data Fig.2a**). A second OPUS-ET round (termed as Expert annotations + OPUS-ET 2nd) further purified the subtomograms to 46,745 subtomograms and achieved 8.5 Å (**Fig.2g; Ext. Data Fig.1d, 2a, Ext. Data Table 1**).

Overlap analysis between OPUS-ET’s picks and human expert annotations showed high purification efficiency by OPUS-ET: between the 150,864 subtomograms in human expert set and the 783,000 subtomograms in template-matching set, 37,489 subtomograms were in common (**Ext. Data Fig.2b**). OPUS-ET Classes 1∼4 recovered 61% (22,778 subtomograms) (**Ext. Data Fig. 2b**), 88% (20,587 subtomograms) of which overlapped with OPUS-ET Classes 3∼13 when purifying human expert annotation (**Ext. Data Fig. 2c**).

**Figure 2.**
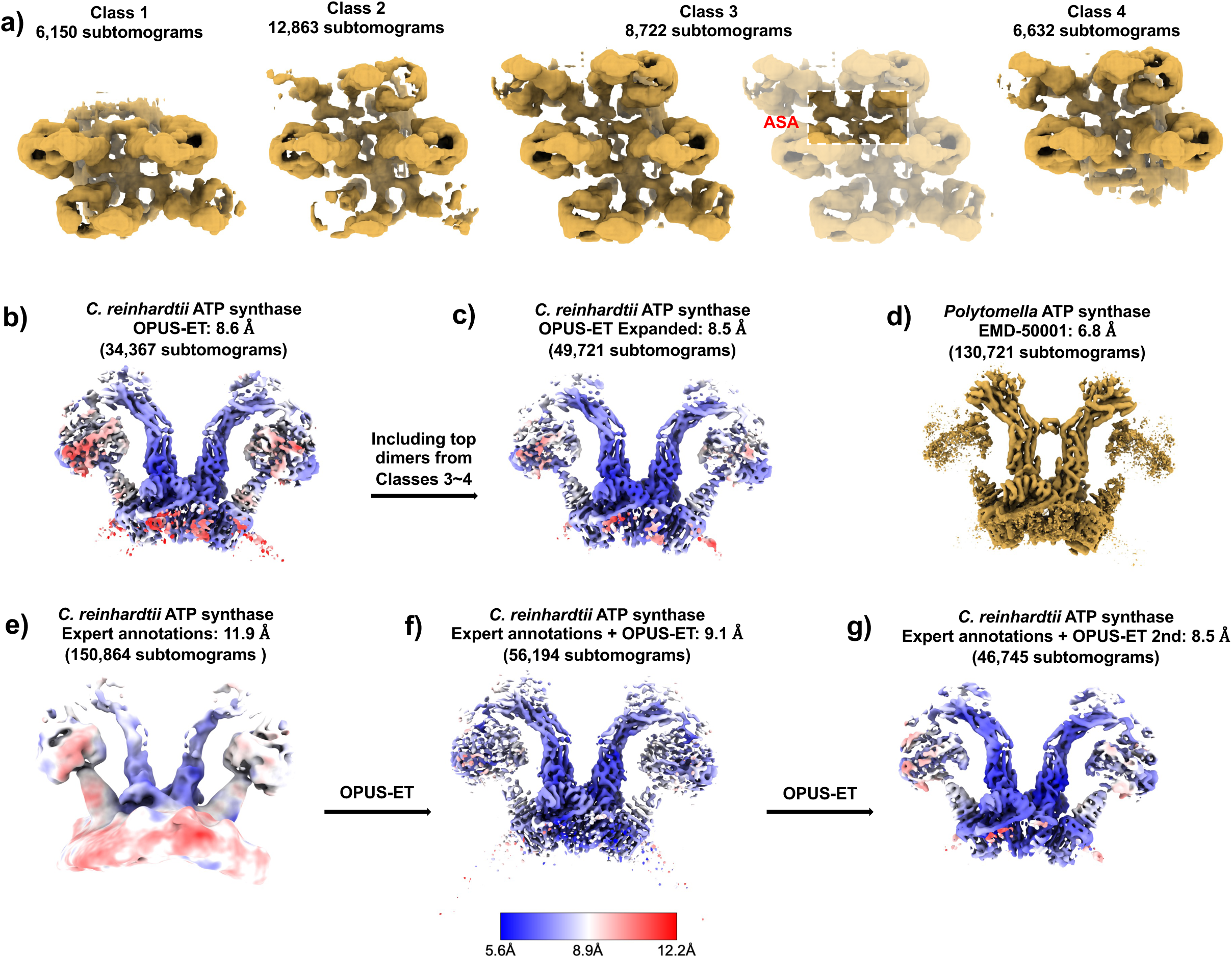
Reconstructions of *C. reinhardtii* ATP synthase dimer *in situ*. **a)**. Distinct oligomeric states identified by OPUS-ET, including dimers, tetramers, and hexamers interlocked by ASA subunits (white dashed box). **b∼g)**. Comparative reconstructions from multiple datasets with indicated resolutions and subtomogram numbers. Density maps are contoured at 4.0𝜎 level in **e)** and 5.5𝜎 level in other maps. **b).** 8.6 Å average from 34,367 subtomograms recovered in OPUS-ET Classes 1∼4. **c).** 8.5 Å average from 49,721 subtomograms by including 15,354 additional subtomograms from the top dimers in Classes 3 and 4 (OPUS-ET Expanded). **d).** Reference *Polytomella* ATP synthase dimer (6.8 Å). **e).** Expert-annotated dataset (150,864 subtomograms) refined to 11.9 Å without helical details. **f).** OPUS-ET-filtered subset (56,194 particles) refined to 9.1 Å showing clear helices. **g).** Second-round purified set (46,745 subtomograms) yielding 8.5 Å map with improved local density.

We next compared OPUS-ET with tomoDRGN using 300,000 *C. reinhardtii* subtomograms picked by template matching from 100 tomograms. For fair comparison, OPUS-ET’s conformation-latent space was turned off (referred to as “OPUS-ET-comp” hereafter to be distinguished from the dual latent disentanglement used in OPUS-ET). OPUS-ET-comp cleanly separated ATP synthase dimers to Classes 11∼12 (**Ext. Data Fig.3a**), whereas tomoDRGN classes were largely mixed, with Classes 5∼6 possibly representing low-resolution dimers (**Ext. Data Fig.3b**). Under identical refinement settings in M with C_2_ symmetry, OPUS-ET-comp (20,693 subtomograms) achieved 9.4 Å, representing a 3.1 Å improvement over tomoDRGN’s reconstruction (23,920 subtomograms), and resolved peripheral-stalk and ASA subunits (**Ext. Data Fig.3c∼g, Ext. Data Table 1**).

To evaluate the impacts of initial templates on the performance of dual-latent OPUS-ET, we analyzed template matching results obtained using low-pass filtered templates at 15 Å, 25 Å or 35 Å cutoffs on the same 100 tomograms. For all templates, OPUS-ET cleanly separated ATP synthase dimers (**Ext. Data Fig.4a∼c**). Under identical refinement settings in M with C_2_ symmetry, OPUS-ET’s selection with 15 Å low-pass filtered template (21,352 subtomograms) achieved 9.2 Å, 25 Å low-pass filtered template (20,230 subtomograms) achieved 9.1 Å, while OPUS-ET’s selection with 35 Å low-pass filtered template (22,412 subtomograms) achieved 9.8 Å (**Ext. Data Fig.4d∼k**). In all cases, the particles picked by OPUS-ET retained high overlapping ratios (≥ 88%) with the expert annotations (**Ext. Data Fig.4k**), and the final reconstructions resolved peripheral-stalk and ASA subunits with comparable local resolutions (**Ext. Data Fig.4g∼i**).

Collectively, these results demonstrate the superb performance and robustness of OPUS-ET for analyzing *C. reinhardtii* ATP synthase heterogeneity. Starting from noisy template-matching data, OPUS-ET achieved 8.5 Å averages within two rounds of filtering and revealed *in situ* oligomeric assemblies. The composition only version of OPUS-ET, OPUS-ET-comp, improved the reconstruction resolution by 3.4 Å on expert-annotated data and substantially outperformed tomoDRGN on the validation dataset, delivering higher-resolution structures in a single classification pass. Furthermore, OPUS-ET remained stable even for highly degraded templates, maintaining sub-nanometer reconstruction and high overlapping ratios.

### Enrichment of eukaryotic 80S ribosomes from cellular tomograms

We next compared OPUS-ET-comp and tomoDRGN^33^ using ten *Schizosaccharomyces (S.) pombe* tomograms from the DeePiCt dataset^39^ (EMPIAR-10988). In the original study^39^, 25,901 manually-annotated 80S ribosome subtomograms yielded a 9.2 Å average with low local resolutions (**Ext. Data Fig.5a∼b**).

Using pyTOM^16^ template matching with a 27 Å low-pass filtered *S. cerevisiae* 80S ribosome map (EMD-3228)^9^ as the template, we picked 50,000 subtomograms and trained OPUS-ET-comp and tomoDRGN^33^ for 38 epochs. By UMAP visualization of learned composition latent space, OPUS-ET-comp Classes 12∼17 (17,215 subtomograms) formed a distinct cluster with complete ribosomal densities and resolved RNA features, whereas all other classes contained non-ribosomal species (**Ext. Data Fig.5c**). In marked contrast, tomoDRGN’s latent space was continuous and less discriminative, with Classes 5∼9 on the right (13,312 subtomograms) showing blurred ribosomal densities (**Ext. Data Fig.5d**). Under identical refinement conditions as DeePiCt, OPUS-ET-comp achieved a 7.4 Å average with improved local resolution, whereas tomoDRGN reached 8.7 Å with an overall lower local resolution (**Ext. Data Fig.5a and e∼f, Ext. Data Table 1**). Fitting the *S. cerevisiae* 80S atomic structure (PDB code 9BDP)^40^ confirmed that the OPUS-ET-comp map resolved secondary-structure elements (**Ext. Data Fig.5e, inset**).

We then compared filtered subtomogram sets with DeePiCt’s expert annotations. Among 50,000 template-matched subtomograms, 21,351 overlapped with DeePiCt’s expert annotations, 66% of them (14,039) were recovered by OPUS-ET-comp Classes 12∼17 and 45% of them (9,514) by tomoDRGN Classes 5∼9 (**Ext. Data Fig.5g∼h**). Tilt-series distribution analysis (**Ext. Data Fig.5i∼j**) showed that OPUS-ET-comp substantially enriched ribosomes than tomoDGRN.

To dissect the contributions of the conformation decoder to the overall performance of OPUS-ET, we performed controlled pose-perturbation tests on the *S. pombe* 80S ribosome dataset, in which the initial particle orientations and translations obtained from template matching were deliberately corrupted prior to training. Rotational perturbations (≤ 30°) were generated by composing the original rotation with a random axis–angle rotation whose axis was sampled uniformly on the sphere and whose angle was drawn uniformly from [0°, θ]; translational perturbations (|Δ| ≤ 29.2 Å, equivalent to |Δ_’_| ≤ 5.0 pixels per dimension at 3.37 Å/voxel) were applied as 3D uniform noise on the particle coordinates.

Under all tested perturbation levels (within 5°+11.7 Å, 10°+17.5 Å, 20°+23.3 Å and 30°+29.2 Å), OPUS-ET consistently outperformed the composition-only model (OPUS-ET-comp) in all comparative measures including overlapping ratio (**Ext. Data Fig.6a**), final reconstruction resolution (**Ext. Data Fig.6b∼c**), as well as reduction of both rotational and translational deviations relative to the M-refined reference poses (**Ext. Data Fig.6d**). Furthermore, UMAP visualizations showed that OPUS-ET maintained a clear separation of ribosome from background noises across all perturbation levels, whereas OPUS-ET-comp showed progressive merging of them (**Ext. Data Fig.6e∼f**).

To gauge the dependence of OPUS-ET to initial templates, we compared template-matching results using as a template the 27 Å or 40 Å low-pass-filtered *S. cerevisiae* 80S ribosome or 27 Å low-pass-filtered rabbit 80S ribosome, a more distant homologue^41^ (EMDB-12756)(**Ext. Data Fig.6g∼i**). In all cases, OPUS-ET separated ribosome particles very well from other particles in UMAP visualization of learned composition latent space, recovered comparable particle sets (17,421, 15,430 and 16,151 particles, respectively), maintained high overlapping ratios (≥ 86%) with the expert annotations, and yielded comparable final reconstruction resolutions of ∼7.0 Å (**Ext. Data Fig.6h∼i**).

In summary, for *S. cerevisiae* 80S ribosome, a single pass of OPUS-ET-comp heterogeneity analysis produced a high-purity particle set that yielded a 7.4 Å reconstruction, which is 1.8 Å better than expert annotation and 1.3 Å better than tomoDRGN, with local resolutions approaching the Nyquist limit (6.7 Å). Additional analysis with introduced perturbation or using lower-filtered or more distant templates demonstrated that OPUS-ET remained robust despite imperfect initial pose or degraded template quality, supporting its potential use even when only distant structural homologues or low-resolution references are available.

### Identification of low-abundance complexes: *S. pombe* fatty-acid synthase

Fatty acid synthase (FAS) is a sparsely populated complex in tomograms of wild-type *S. pombe* (DeePiCt dataset, EMPIAR-10988)^39^, where only 366 subtomograms were manually annotated and refined to 18.5 Å with poor local resolutions after multiple rounds of refinements in M^4^ (**Ext. Data Fig.7a∼b**).

Using pyTOM with a 27 Å low-pass filtered *S. cerevisiae* FAS map^42^ (EMDB code EMD-1623) as template, we selected 4,800 subtomograms and trained OPUS-ET-comp and tomoDRGN under identical conditions. By UMAP visualization of learned composition latent space, OPUS-ET-comp Class 19 (221 subtomograms) formed an isolated island in the UMAP and showed complete FAS densities (**Ext. Data Fig.7c**). In contrast, tomoDRGN Class 0 (170 subtomograms) contained incomplete, low-contrast densities (**Ext. Data Fig.7d**). The map of OPUS-ET-comp Class 19 revealed higher-resolution features, particularly in regions marked by red solid and dashed boxes (**Ext. Data Fig.7e**) than that of tomoDRGN Class 0 (**Ext. Data Fig.7f**). Refinement in M with D₃ symmetry produced a 14.0 Å average for OPUS-ET-comp, compared with 18.5 Å for tomoDRGN, reflecting a 4.5 Å improvement with markedly better local resolution (**Ext. Data Fig.7a and g∼h, Ext. Data Table 1**).

Among the 4,800 template-matched subtomograms, 115 overlapped with the 366 DeePiCt’s expert annotations, 77% of them (89 particles) were recovered by OPUS-ET-comp Class 19, and only 51% of them (59 particles) by tomoDRGN Class 0 (**Ext. Data Fig.7i∼j**). Tilt-series distributions confirmed higher overlap ratios for OPUS-ET-comp (**Ext. Data Fig.7k∼l)**.

Visual inspection of tilt series TS_045 showed that OPUS-ET-comp particles displayed coherent FAS-like features, whereas several DeePiCt’s expert annotations and tomoDRGN particles represented membrane fragments or noise (**Ext. Data Fig.7m,** indicated by red circles).

Furthermore, we carefully inspected the 26 subtomograms that were overlapped with the DeePiCt’s expert annotations but missed by OPUS-ET-comp (115-89=26 subtomograms), and found they all lacked distinct features expected for FAS (**Ext. Data Fig.7n**), justifying their absence in the particle set picked by OPUS-ET-comp.

To evaluate sensitivity to the number of latent classes, we repeated the *S. pombe* FAS analysis across a range of K values (K = 5, 10, 20, and 40) for both OPUS-ET-comp and tomoDRGN (**Ext. Data Fig.8a∼d**). At K = 5, neither method achieved sufficient resolution to isolate FAS: OPUS-ET-comp’s best-matching cluster contained 622 subtomograms with 93 overlapping with DeePiCt expert annotations but yielded only a 22.1 Å reconstruction, indicating that FAS particles are co-clustered with background species at this coarse granularity.

Meanwhile, tomoDRGN’s corresponding cluster was even larger (1,472 subtomograms) and reached only 24.5 Å in final reconstruction. For K of 10, 20 or 30, OPUS-ET-comp converged to a stable FAS cluster of ∼220 subtomograms, with ∼99% internal consistency across K values (measured as overlap with the K = 20 cluster) and nearly identical reconstruction resolutions of ∼14.2 Å (**Ext. Data Fig.8a∼d**). Overlap with the DeePiCt’s expert annotation set remained constantly around ∼90 subtomograms. By contrast, the subtomograms selected by tomoDRGN overlapped with the DeePiCt set at around ∼60 for K ≥ 10, and resulted in reconstruction resolution ranged of 18.2∼24.4 Å (**Ext. Data Fig.8a∼d**). Multiple runs using different random seeds yielded essentially the same results for both OPUS-ET-comp and tomoDRGN (**Ext. Data Fig.8e**).

To assess FAS classification robustness under imperfect pose initialization, we applied rotational and translational perturbations (10°+17.5 Å, 20°+23.3 Å, or 30°+29.2 Å) to the initial FAS candidates prior to OPUS-ET training, and compared the results against the unperturbed baseline (termed as “Without Perturbation”) (**Ext. Data Fig.9a∼c**). In contrast to OPUS-ET-comp, the conformation decoder in the OPUS-ET full model predicts a corrected pose for each particle, thereby directly reducing the deviation with respect to the pose from high-resolution M refinement result. Without perturbation, OPUS-ET recovered 228 particles (216 overlapping with the K = 20 OPUS-ET-comp reference) at 14.1 Å resolution, with translational deviations corrected from 17.2 Å to 11.8 Å. Across all perturbation levels, OPUS-ET consistently separated FAS particles into an isolated cluster (**Ext. Data Fig.9a**). Under moderate perturbation levels (10°+17.5 Å and 20°+23.3 Å), it recovered 232 and 212 particles (overlapping rate of 98% and 87%) with 14.5 Å and 14.8 Å resolution, respectively (**Ext. Data Fig.9b∼c**). Under the highest perturbation level (30°+29.2 Å), it recovered 210 particles (154 overlapping) with 18.1 Å resolution. For the three perturbation levels, the conformation decoder reduced translational deviations by 7.1, 5.2 and 5.2 Å, and rotational deviations by 0.7, 3.6 and 3.4°, respectively (**Ext. Data Fig.9c**). Therefore, in applying to FAS, OPUS-ET demonstrated robustness across K values (K = 10 to 40), runs with different random seeds and imperfect initial pose.

These examples have established that OPUS-ET efficiently resolves compositional heterogeneity and yields purified, high-resolution particle sets across diverse cellular targets. The perturbation experiments further demonstrate that the conformation decoder not only maintains robust particle classification under imperfect initial pose, but also effectively corrects pose errors during training.

Having established the robustness of our dual-latent OPUS-ET framework and the significant contributions of the novel conformation decoder, we next asked whether it could also recover biologically meaningful continuous conformational landscapes directly from cryo-ET data.

### Rotary and inter-subunit dynamics of mitochondrial ATP synthase dimers

We applied OPUS-ET to the M-refined set of 49,721 subtomograms (8.5 Å) of *C. reinhardtii* mitochondrial ATP synthase with C_2_ symmetry expansion to examine multiscale conformational changes within the central dimer. During training, the conformation decoder used a two-subunit model centered on the F₁-head center of mass (COM). OPUS-ET revealed the flexible coupling between F_0_ and F_1_ subcomplexes within each monomer while capturing relative movements between monomers.

It is well established that the central stalk of ATP synthase undergoes a 360° rotation in 120° steps, defining three primary rotary states^43^. OPUS-ET recovered these states by clustering the composition latent space and reconstructing maps at class centers (**Ext. Data Fig.10a**).

Classes 23, 7, and 19 correspond to the three rotary states, as validated by atomic structure overlays (**Fig.3a**). In superposition, the central stalk of Class 23 points left, Class 7 centers, and Class 19 points right (**Fig.3b, Upper**), with ∼120° separation in rotated views (**Fig.3b, Lower**).

Furthermore, OPUS-ET resolved rotary substates distinguished by F₁-head rotation relative to the peripheral stalk. Rigid-body fittings identified Classes 17 and 19 as Substates 3C and 3A^43^, respectively (**Fig.3c**). During the 3A to 3C transition, the F₁-head rotates counter-clockwise, supported by density changes in the ⍺-subunit near the peripheral stalk (**Fig.3d, Upper**). Superposed atomic structures show coordinated rearrangements: ASA and oligomycin sensitivity–conferring protein (OSCP) shift rightward while ⍺ and β subunits rotate counter-clockwise, the central stalk of Class 19 also rotates counter-clockwise relative to Class 17, confirmed by density differences in the sectional view and the superposition of fitted γ subunits (**Fig. 3d**). These comparisons confirm that F₁-head and central stalk rotate synchronously within a primary state^43^.

**Figure 3.**
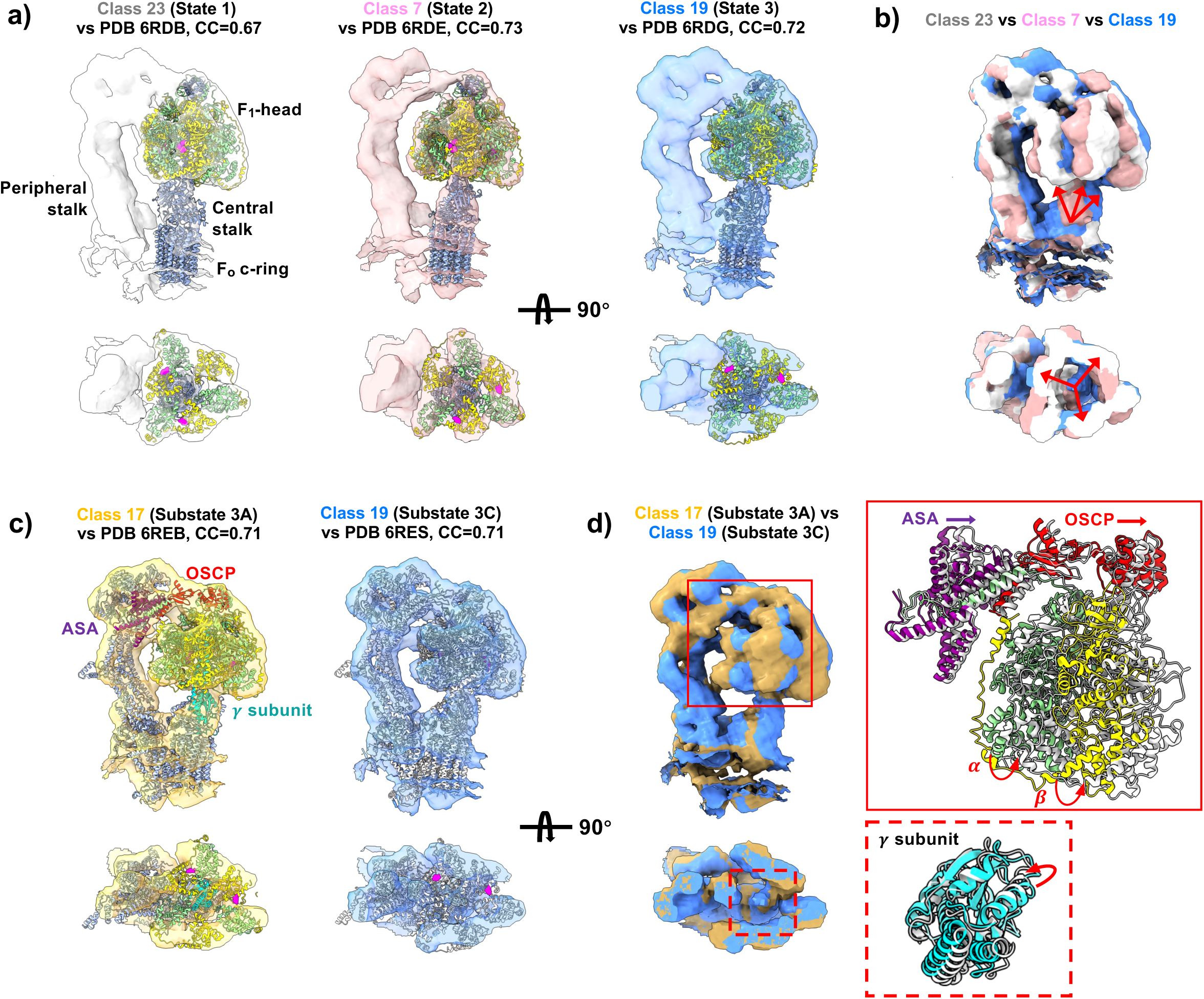
Rotary states of *C. reinhardtii* ATP synthase *in situ*. **a)**. Three primary rotary states, Class 23 (State 1), Class 7 (State 2), and Class 19 (State 3), with fitted atomic models highlighting the central stalk (dark blue), F₁-head α subunits (light green), and β subunits (yellow) in side view **(Upper)** and top view showing section through F_1_ at the level of bound ADP (magenta spheres) **(Lower)**. **b)**. Superposed states illustrate ∼120° rotary offset of the central stalk between sequential states (arrow directions) in side view **(Upper)** and top view showing section through F_1_ at the level of bound ADP (magenta spheres) **(Lower)**. **c)**. Two rotary substates (3A and 3C) resolved by OPUS-ET with fitted atomic models showing α subunits (light green), β subunits (yellow), OSCP (red), γ subunit (turquoise), ASA (purple) and other peripheral stalk components (blue) in side view **(Upper)** and top view showing section through F_1_ at the level of bound ADP (magenta spheres) **(Lower)**. **d)**. Overlay of Substates 3A (brown) and 3C (blue) shows counter-clockwise rotation of α/β subunits and rightward shift of ASA/OSCP (red solid box zoom-in), as well as counter-clockwise rotation of γ subunit (red dashed box zoom-in), defining F₁–F₀ coupling mechanics.

Whereas discrete clustering partitions the latent space into fixed classes, principal-component analysis (PCA) identifies directions of maximal variance in the particle distribution, capturing dominant modes of structural variation continuously. Traversing the composition latent space along PC9 visualized the concerted rotation of F_1_-head and central stalk across states (**Ext. Data Fig.10b**). Along PC9, the positive end (1.4PC9) corresponds to Substate 3C, the midpoint (−0.3PC9) to 3B, and the negative end (−1.2PC9) to 1A^43^ (**Fig.4a∼c**). Fitted atomic structures yielded CC values of 0.69∼0.72. Superimposed maps show progressive displacements: from 3C→3B, OSCP rotates with F₁-head clockwise while ASA shifts rightward, γ-subunit rotates by ∼10° counter-clockwise and β-subunits rotate by ∼15° clockwise; from 3C→1A, OSCP and F₁-head continue their clockwise rotations, ASA shifts further rightward, γ-subunit rotates by ∼90° counter-clockwise, and β subunits rotate by ∼30° clockwise (**Fig.4**).

**Figure 4.**
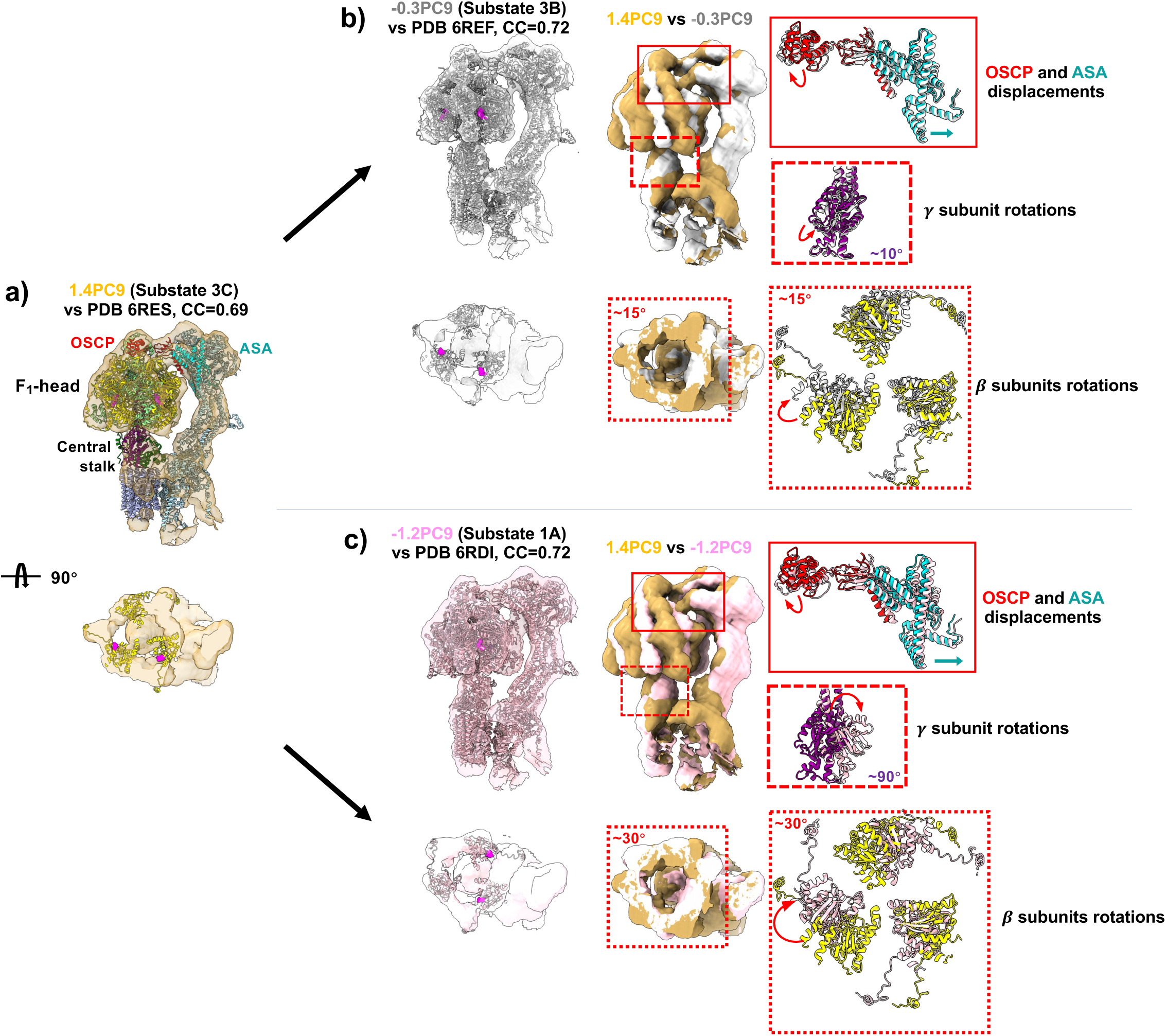
Dynamics of *C. reinhardtii* ATP synthase *in situ*. Principal-component traversal (PC9) reveals rotary continuum from **a)** Substate 3C (1.4 PC9, yellow), through **b)** intermediate 3B (–0.3 PC9, grey), to **c)** Substate 1A (–1.2 PC9, pink); each map fitted with atomic models (CC=0.69∼0.72). Lower row shows top view sectioned through F_1_ at the level of bound ADP (magenta spheres). **a)**. Superposition of 1.4PC9 and the structure for Substate 3C, with OSCP (red), ASA (turquoise), α subunits (light green), β subunits (yellow), and γ subunit (purple). **b)**. Superpositions of Substates 3C and 3B show OSCP clockwise rotation and ASA rightward shift (Upper, red solid box zoom-in), 10° counter-rotation for γ subunit (Upper, red dashed box zoom-in), and 15° clockwise rotation for β subunit (Lower, red dashed box zoom-in). **c)**. Superpositions of Substates 3C and 1A show OSCP clockwise rotation and ASA rightward shift (Upper, red solid box zoom-in), ∼90° counter-rotation for γ subunit (Upper, red dashed box zoom-in) and ∼30° clockwise rotation for β subunit (Lower, red dashed box zoom in), completing one 120° rotary step (**Suppl. Video 1**).

Collectively, these motions complete one elementary 120° step, producing concerted counter-rotations that reconcile the F₀ ring–F₁ head mismatch^43^ (**Suppl. Video 1**). This counter-rotation pattern was reproduced by traversing PC8 in the learned composition latent space of an independent training run (**Suppl. Video 2**). Thus, OPUS-ET captures the entire F₀–F₁ flexible coupling mechanics *in situ*.

OPUS-ET further revealed inter-subunit dynamics through its conformation decoder. Traversing the first two principal components (DPC1 and DPC2) reconstructed two orthogonal normal modes (**Ext. Data Fig.10c∼d**). Fitted atomic structures (PDB codes: 6RDE and 6RD4) show that along DPC1, the two monomers rotate in opposite directions, bending the dimer around parallel axes (**Suppl. Video 3**). Along DPC2, monomers rotate in the same direction, producing a twisting motion around orthogonal axes (**Suppl. Video 4**). These motions illustrate that OPUS-ET can extract normal-mode-like dynamics directly from cryo-ET data.

Through these analyses, OPUS-ET reconstructs the full F₀–F₁ flexible coupling mechanics and reveals normal-mode-like bending and twisting of dimers *in situ*.

### tRNA-translocation intermediates and elongation-factor cycling in *S. pombe* 80S ribosomes

Eukaryotic 80S ribosomes synthesize proteins by translating mRNA through large-scale conformational changes driven by elongation factors (eEFs), with three tRNA binding sites (A, P, and E) spanning from the 40S head to the 60S subunit. During elongation, these sites are sequentially occupied and released by tRNAs and eEFs. To analyze the underlying compositional variability and dynamics, we trained OPUS-ET on the M-refined OPUS-ET set (17,215 subtomograms). The conformation decoder applied a two-subunit model to capture the relative 40S∼60S displacements, with each subunit centered at its COM.

Among 30 well-separated classes, Classes 26∼29 contained non-ribosomal densities, Class 24 comprised isolated 60S subunits, and the remaining classes displayed complete 80S ribosomes (**Ext. Data Fig.11a**). We identified distinct eEFs occupancies on 80S ribosomes (**Fig.5**). Classes 17 and 25 showed an ear-like density at the 40S head assignable to eEF3 by model fitting (PDB code 7B7D^44^) (**Fig.5a∼b**). Most notably, OPUS-ET distinguished eEF1A·tRNA in Class 17 (CC = 0.74 to PDB code 9BDP) and eEF2 in Class 25 (CC = 0.70 to PDB code 7OSM) (**Fig.5a∼b**), two factors that occupy an overlapping 60S-proximal site and are rarely separable without high local resolution.

**Figure 5.**
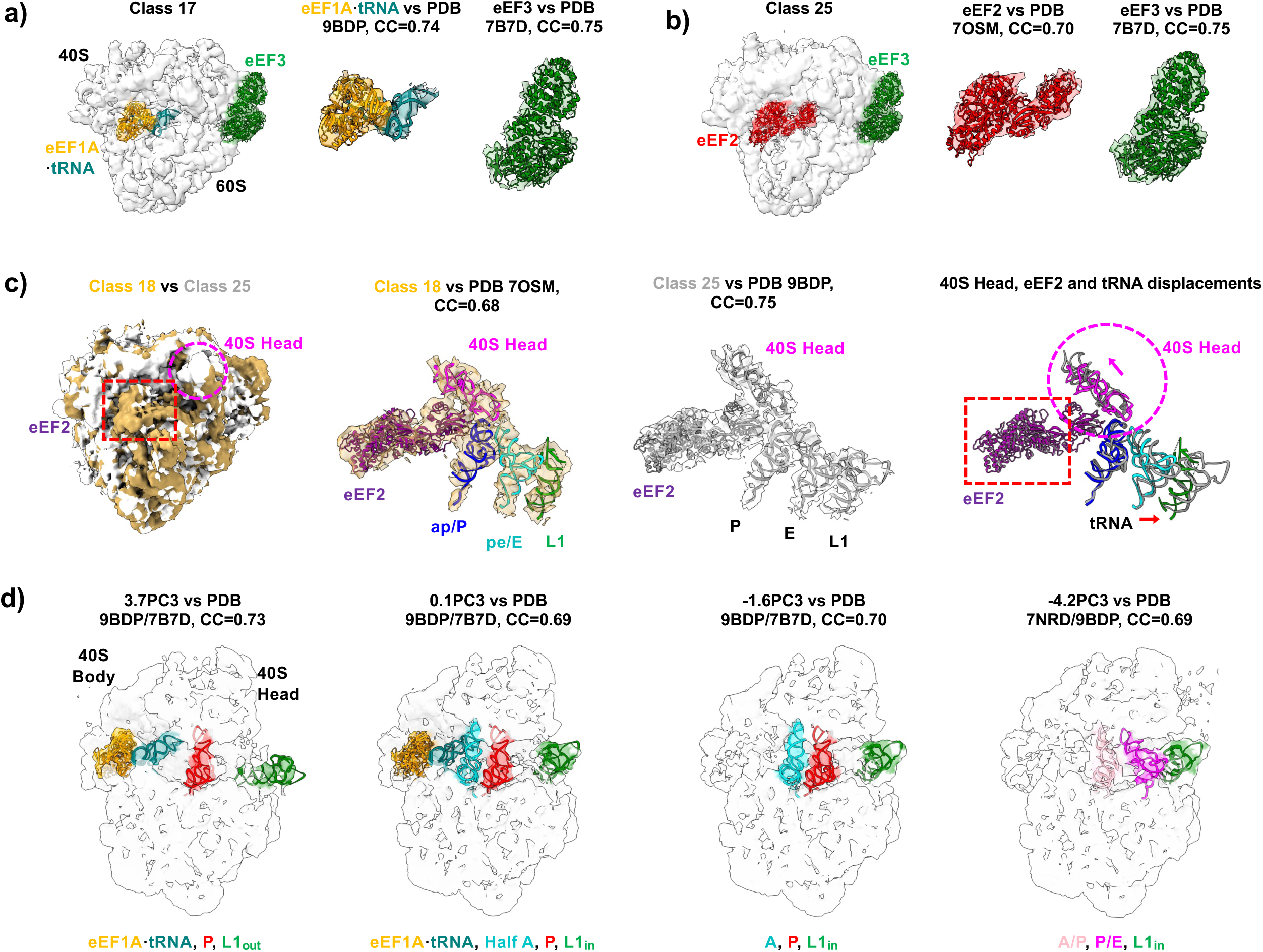
Translocation-intermediate states of *S. pombe* 80S ribosome *in situ*. **a∼b)**. OPUS-ET maps of ribosomes bound to **a)** eEF1A·tRNA and eEF3, and **b)** eEF2 and eEF3, each fitted with atomic models (CC=0.70∼0.75). **a)**. Class 17 with eEF1A×tRNA (orange and teal) and eEF3 (green). **b)**. Class 25 with eEF2 (red) and eEF3 (green). **c)**. Comparative view of eEF2-bound states: Class 18 (hybrid ap/P and pe/E tRNAs) and Class 25 (classical P/E tRNAs). Superposition shows clockwise 40S-head rotation and tRNA rightward movement while eEF2 anchors in the A-site as a “locking pawl”. **d)**. PC3 traversal reconstructs tRNA translocation continuum from A/T and P states to A/P and P/E hybrids with L1-stalk closure and counter-clockwise 40S rotation (**Suppl. Video 4**). The structures of eEF1A×tRNA (orange and teal, PDB code 9BDP), A-site (turquoise, PDB code 7B7D), P-site (red, PDB code 9BDP), A/P-site (pink, PDB code 7NRD) and P/E-site (magenta, PDB code 7NRD) tRNAs are fitted into density maps and shown as references. The L1 stalk (green) conformations, L1_out_ and L1_in_, are fitted into density maps using the atomic structure from PDB code 9BDP. All densities are colored consistently with their respective fitted models.

**Figure 6.**
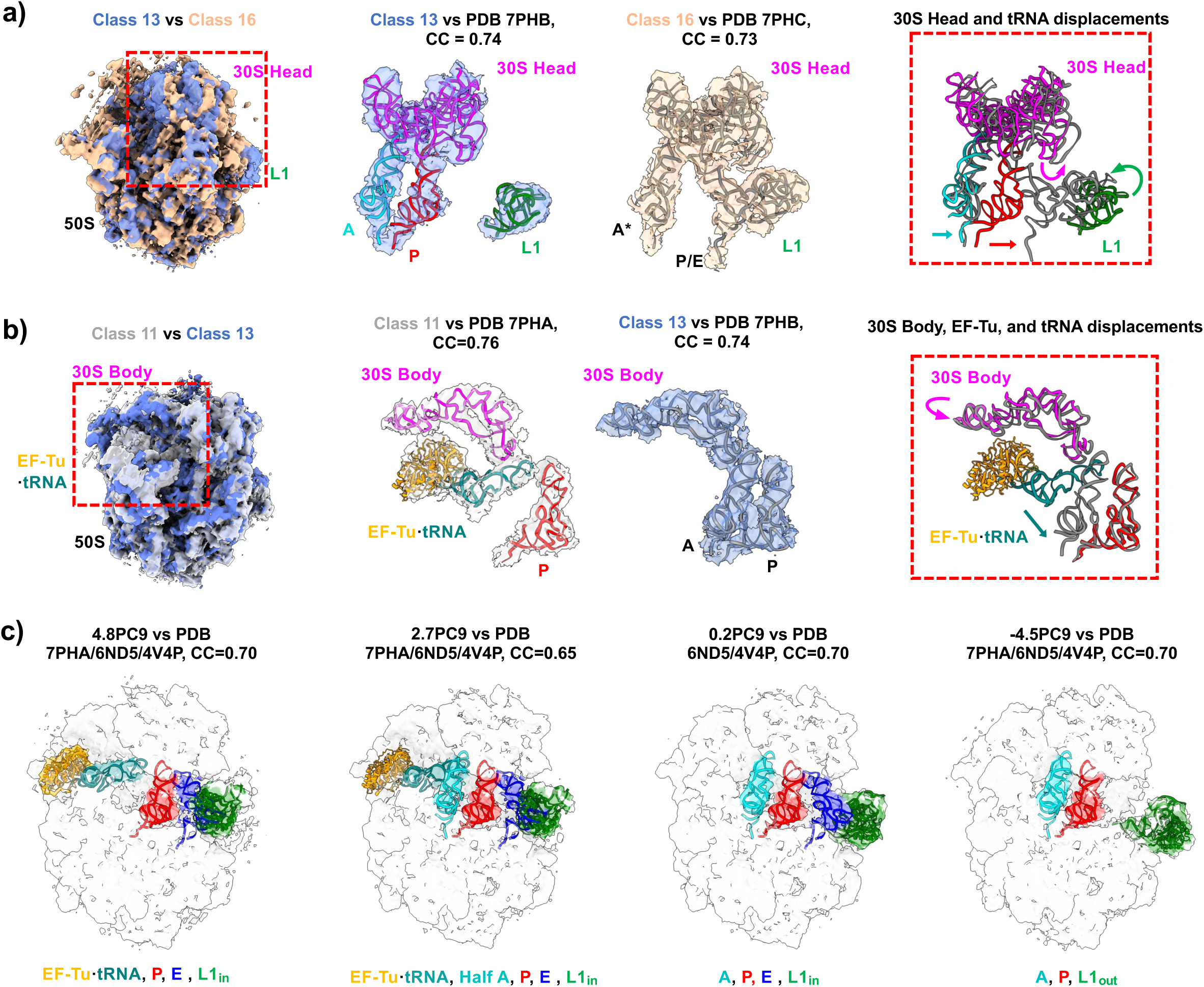
Translocation-intermediate states of *M. pneumoniae* 70S ribosome *in situ*. **a)**. Comparison of Classes 13 (A and P) and 16 (A* and P/E) shows 30S-head counter-clockwise rotation and L1-stalk movement toward E site (red dashed box zoom-in). Atomic structures fitted in Class 13 map are in color: 30S Head (magenta), A-site tRNA (turquoise), P-site tRNA (red) and L1 stalk (green). The arrows in red dashed box are colored according to the corresponding structures. **b)**. Class 11 (EF-Tu·tRNA and P) to Class 13 (A and P) illustrates 30S-body downward roll and A-site delivery of tRNA as EF-Tu releases. Atomic structures fitted in Class 11 map are in color: 30S Body (magenta), EF-Tu×tRNA (orange and teal), and P-site tRNA (red). The arrows in red dashed box are colored according to the corresponding structures. **c)**. PC9 traversal visualizes full tRNA delivery and E-site release sequence with L1-stalk switch (L1_in_ → L1_out_) and correlated EF-Tu movements (**Suppl. Video 7**). The structures of A- (turquoise), P- (red) and E-site (blue) tRNAs from *T. thermophilus* Cm-bound 70S ribosome (PDB code 6ND5) and the structures of EF-Tu×tRNA (orange and teal) from 70S ribosome in Cm-treated *M. pneumoniae* cell (PDB code 7PHA) are fitted into density maps and shown as references. The L1 stalk (green) conformations, L1_out_ and L1_in_, are fitted into density maps using the atomic structure from PDB code 4V4P^55^. All densities are colored consistently with their respective fitted models.

Classes 18, 19, 21, 22, and 25 formed a major eEF2-containing cluster in composition latent space (**Ext. Data Fig.11a**). Comparative analysis revealed eEF2 as a “locking pawl” during tRNA translocation^45^ (**Fig.5c**). Class 18 contains hybrid ap/P- and pe/E-site tRNAs (fitted with PDB code 7OSM^45^), whereas Class 25 contains classical P- and E-site tRNAs (fitted with PDB code 9BDP^40^) (**Fig.5c, Middle**). Superpositions show eEF2 anchors at the A-site while neighboring tRNAs shift in the direction opposite to the 40S-head rotation (**Fig.5c, Right**), acting as a pawl that prevents backward slippage, which is consistent with prior observations^45^. Additional translocation intermediates uncovered in our analysis included Class 4 (A- and P-site tRNAs with eIF5A at E-site), Class 13 (eEF1A·tRNA with P-site tRNA and eIF5A at E-site), Class 17 (eEF1A·tRNA with P- and E-site tRNAs), and Class 22 (eEF2 with A/P- and P/E-site tRNAs) (**Ext. Data Fig.11b**).

The elongation cycle of the 80S ribosome begins with eEF1A delivering tRNAs to the vacant A-site, proceeds through intermediate hybrid states (A/P and P/E, ap/P and pe/E), and concludes with tRNAs occupying the P- and E-sites^45^. OPUS-ET’s principal-component traversal aligned with a large portion of this cycle (**Ext. Data Fig.11c**). Along PC3, at 3.7PC3, we observed eEF1A·tRNA, a P-site tRNA and an empty E-site, with L1 stalk in L1_out_ conformation. At 0.1PC3, eEF1A×tRNA partly occupies A-site while L1 stalk transits to L1_in_, closing E-site. At -1.6PC3, the tRNA is delivered into A-site, and L1 stalk remains in L1_in_. At - 4.2PC3, A/P- and P/E-site tRNAs appear and L1 stalk contacts the P/E-site tRNA, underscoring its role in hybrid-state formation (**Fig.5d**). During this progression, the 40S head rotates counter-clockwise, narrowing the A–P to P–E inter-subunit gaps (**Suppl. Video 5**). Similar tRNA translocation in these sites can again be reproduced by traversing the PC4 of learned composition latent space in an independent training run (**Suppl. Video 6**).

Analysis of the conformation latent space revealed two orthogonal normal-mode-like motions (DPC1, DPC2). We fitted 40S and 60S from PDB code 7B7D independently into the density maps reconstructed along each DPC to quantify displacements (**Ext. Data Fig.11d∼e**). Along DPC1, the 40S rotates counter-clockwise about its body–head axis while the 60S rotates clockwise about its central axis (**Ext. Data Fig.11d**), producing a twisting motion (**Suppl. Video 7**). Along DPC2, both subunits undergo coordinated translation and rotation around a 40S-centered axis, yielding a smooth displacement gradient across the ribosome (**Ext. Data Fig.11e**, **Suppl. Video 8**).

These results demonstrate that OPUS-ET recovers inter-subunit dynamics directly from cryo-ET data without manual state selection. Such analyses reveal a continuous progression of conformational states spanning hybrid and classical tRNA configurations, and capture inter-subunit counter-rotation during translocation.

### Translation dynamics of chloramphenicol-treated *Mycoplasma (M.) pneumoniae* 70S ribosomes

To further test OPUS-ET, we analyzed chloramphenicol (Cm)-treated *M. pneumoniae* 70S ribosomes^33,46^ (EMPIAR accession code 11843), previously used to benchmark tomoDRGN^33^ and CryoDRGN-ET^34,47^. Using M-refined 22,291 subtomograms, OPUS-ET identified two new translocation-intermediate states absent from earlier analyses by CryoDRGN-ET or RELION^34,47^: Class 19 containing tRNAs at A-, P- and E-sites and Class 23 containing EF-Tu×tRNA with tRNAs at P- and E-sites (**Ext. Data Fig.12a∼b, Ext. Data Table 1**).

OPUS-ET reconstructed concerted conformational changes during transitions among intermediates. For example, Class 13 represents a state with A- and P-site tRNAs (fitted with PDB code 7PHB^47^), while Class 16 corresponds to a state with tRNAs at A*- and P/E-sites, (PDB code 7PHC^47^) (**Fig.6a**). Superposition revealed counter-clockwise rotation of the 30S head and leftward movement of the L1 stalk toward the E-site (**Fig.6a, Right**). Class 11, containing EF-Tu×tRNA and P-site tRNA (PDB code 7PHA^47^) (**Fig.6b**), captured an earlier delivery stage; comparison with Class 13 showed the 30S body rolling downward as EF-Tu dissociates and tRNA enters the A-site (**Fig.6b**, **Right**).

Principal-component traversal in OPUS-ET’s 12-dimensional composition latent space revealed correlated changes linking these intermediates. PC9 aligned with tRNA delivery by EF-Tu cofactor and E-site tRNA release facilitated by L1 stalk (**Fig.6c** and **Ext. Data Fig.12c**): at 4.8PC9, EF-Tu×tRNA occupies the A-site at 30S subunit, with other tRNAs at P- and E-sites and L1 stalk in L1_in_ conformation; at 2.7PC9, EF-Tu×tRNA partly occupies A-site, while P- and E-site tRNAs, and L1 stalk remain in similar conformations; at 0.2PC9, a triple-tRNA state (A, P, E) forms; and at -4.5PC9, the E-site tRNA dissociates as the L1 stalk transitions to L1_out_ conformation (**Fig.6c**). These transitions visualized the tRNA delivery-to-release continuum observed in the bacterial elongation cycle (**Suppl. Video 9**).

Traversing PC8 revealed occupancy changes of the L9 protein on the 50S subunit (**Ext. Data Fig.12d**), undetected in previous deep-learning analyses by CryoDRGN-ET and tomoDRGN^33,34^. Densities for L9 (PDB code 7PHA) disappear when moving from –4.7 PC8 to 5.8 PC8, suggesting dynamic association with adjacent rRNA regions.

The translation-related structural variation of 70S ribosome in Cm-treated *M. pneumoniae* cells were previously reconstructed by CryoDRGN-ET^34^ and tomoDRGN^33^ **(Suppl. Video 10**), required either manual interpolation or KMeans sampling to visualize dynamics. In contrast, OPUS-ET reconstructed a more complete translation trajectory automatically through continuous latent-space traversal **(Suppl. Video 9)**. Moreover, OPUS-ET maps showed stronger and more complete EF-Tu·tRNA densities (**Ext. Data Fig.12e**).

We further benchmarked the three methods on a reduced dataset (2,990 subtomograms from the first eight tomograms in the *M. pneumoniae* ^33,46^ dataset (EMPIAR code 11843)). OPUS-ET cleanly separated complete 70S ribosomes (Classes 6∼17), 50S subunit (Classes 18), and noise (Classes 0∼5 and 19), whereas tomoDRGN and CryoDRGN-ET failed to distinguish these species (**Ext. Data Fig.13a∼c, Ext. Data Table 1**). Representative subtomogram averages using OPUS-ET’s classification confirmed its separations (**Ext. Data Fig. 13d**).

Collectively, OPUS-ET resolves conformational states along the bacterial elongation cycle and identifies biologically important intermediates and auxiliary-protein rearrangements. OPUS-ET maintains robust performance even on small datasets, an essential feature for studying rare species or limited cellular volumes.

In summary, across four cellular systems, OPUS-ET resolves compositional variability and conformational dynamics within one unified framework. Its latent-space representations not only isolate pure populations of target species but also encode coordinates describing rotary, translational, and bending motions, achieving a comprehensive description of macromolecular heterogeneities. A performance summary table is provided in **Ext. Data Table 1**, and a training summary table is provided in **Ext. Data Table 2**.

## Discussion

Structural heterogeneity presents a fundamental challenge in cryo-ET data processing, limiting both resolution and biological interpretation. Here we introduce OPUS-ET, a deep learning framework that integrates high-resolution structural determination with dynamics modelling for cryo-ET. By encoding subtomograms into two separate low-dimensional latent spaces, composition latent space and conformation latent space, OPUS-ET provides a hierarchical representation for structural heterogeneities, and enables high-resolution recovery of target complexes from noisy inputs with pose assignments derived from template matching. Across multiple benchmark systems, OPUS-ET demonstrates robust performance under varying template quality, initial pose noise, and clustering parameters, establishing it as a powerful tool for *in situ* structural analysis.

Compared to traditional 3D-classification approaches, OPUS-ET achieves both computational and interpretive gains. It can produce highly purified subtomogram sets for target species directly from template-matching results, substantially reducing manual curation and computation. Subsequent M refinement yields reconstructions surpassing expert-curated classifications in our benchmark systems. Crucially, OPUS-ET reconstructs structural continua rather than discrete classes, revealing dominant modes that connect intermediate conformations. This capability is particularly important for identifying short-lived, functionally relevant states^20^.

Compared with previous deep-learning-based approaches, such as tomoDRGN and CryoDRGN-ET, which encode heterogeneity through a single decoder, OPUS-ET uses distinct conformation and composition decoders that play complementary roles. The conformation decoder actively absorbs large-scale motions, while the composition decoder captures other structural differences, including subunit occupancy, assembly state, and residual density variations. This complementarity is especially important when pose uncertainty arises at two levels: at the whole complex level, from noisy pose assignments in template-matching inputs, and at the subunit level, from large-scale inter-subunit movements in multi-subunit complexes. By absorbing these continuous displacements, the conformation decoder removes a source of density variation that would otherwise contaminate the composition latent space, allowing it to form denser, more stable clusters for particles with similar compositions. Perturbation and ablation analyses confirm that this separation stabilizes compositional recovery and improves both pose accuracy and reconstruction resolution relative to tomoDRGN, even under substantial pose perturbations. In real-world multi-subunit complexes, the conformation decoder revealed normal-mode-like structural variations directly from cryo-ET data, such as rotary bending and twisting in ATP synthase or inter-subunit rotation in ribosomes.

Because large-scale motions are absorbed by the conformation decoder, the composition latent space of OPUS-ET captures residual structural differences with higher fidelity, yielding clusters and traversal directions that are more tightly linked to biologically meaningful variation. In the benchmark systems studied here, OPUS-ET achieves equivalent or superior interpretability through a single training-and-traversal step, supported by sub-nanometer reconstructions of ATP synthase and ribosomes and by discovery of biologically important intermediate states, including the hybrid ap/P–pe/E 80S ribosome^45^ and L9 occupancy changes in 70S ribosomes. By contrast, existing approaches such as tomoDRGN^33^ and CryoDRGN-ET^34^ often require additional *post hoc* procedures, including manual latent-code selection, MAVEn sampling, or ensemble averaging, to visualize comparable transitions.

By automating heterogeneity analysis directly from template-matching results with limited human intervention, OPUS-ET significantly reduces the expertise and effort required for high-resolution cryo-ET. Its ability to represent both discrete heterogeneity and continuous structural variation enables direct investigation of coordinated subunit and cofactor rearrangements *in situ*. As cryo-ET expands toward increasingly complex cellular samples, scalable methods that integrate particle purification, structural recovery, and analysis of large-scale structural variation will become essential for interpreting native cellular organization. Future work will explore extension of OPUS-ET to single-particle cryo-EM, where the same dual-latent architecture could address similar heterogeneity challenges therein.

## Methods

### Deformation field

OPUS-ET represents each 3D structure as the combination of a static template volume and a continuous deformation field that specifies voxel displacements relative to this template. Each voxel in the field records its source location in the undeformed template, allowing conformations to be sampled directly.

Formally, the 3D conformation is expressed as the composition of two functions 𝑉(𝒙 + 𝒈(𝒙)): ℝ^3^ → ℝ, where 𝑉(·) is the template which maps a grid point to a voxel value, and 𝒈(𝒙): ℝ^3^ → ℝ^3^ is the deformation field that maps each output coordinate 𝒙 to its source coordinate 𝒙 + 𝒈(𝒙) in template, recording the displacement between the two locations. The output intensity in deformed conformation is obtained by trilinear interpolation at that source location 𝒙 + 𝒈(𝒙) through a spatial transformer^36^. Next, we give a detailed definition of the deformation field.

In OPUS-ET, the total deformation is decomposed into rigid-body motions of predefined subunits. Each subunit is defined by user and specified by a binary mask volume 𝑆 ⊆ ℝ^3^, which defines the set of grid points belonging to it. The effective mass at each grid point is:

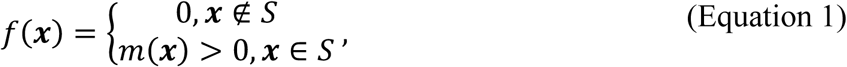

Its center is automatically determined from the mask 𝑆 by computing the center of mass,

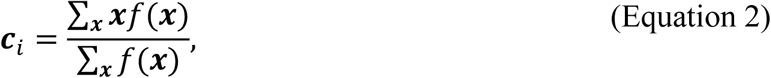

and its principal axes 𝑃_’_ are obtained from the eigendecomposition of the inertia tensor,

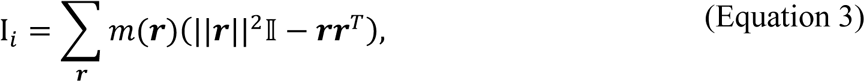

where 𝒓 = 𝒙 − 𝒄_’_. We can further define some shape statistics about the subunit. For a subunit with center 𝒄_’_ ∈ ℝ^3^, and suppose its principal axes form a matrix 𝑃_’_, its semi-principal diameters can be defined as

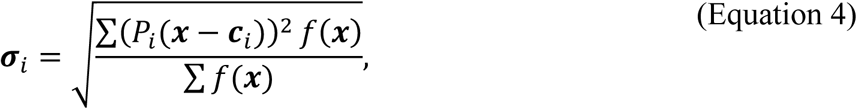

where the summations are over all grid points in the 3D grid. For a grid point 𝒙 = {𝑥, 𝑦, 𝑧} ∈ ℝ^3^ in a subunit 𝑖, suppose the subunit undergoing a rigid-body displacement with a rotation matrix 𝑅_’_ ∈ 𝑆𝑂(3), and translation vector 𝒕_’_ ∈ ℝ^(^, let the center of mass (COM) of the subunit be 𝒄_’_, the rigid-body displacement of this point can be expressed as

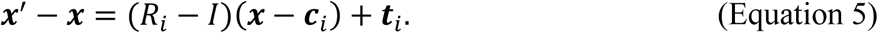

Voxel displacements from multiple subunits are blended smoothly according to their distance from each subunit COM. Specifically, for a grid point 𝒙, suppose the principal axes of subunit 𝑖 form a matrix 𝑃_’_ as preceding, the displacement incurred by the movement of subunit 𝑖 decays as,

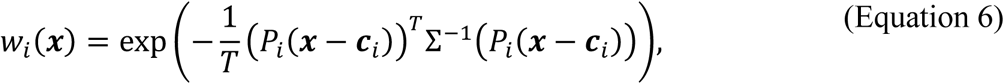

where 𝑥_0_ is the 𝑗th component of the grid point 𝒙, Σ is a diagonal matrix with its diagonal element Σ_00_ = 𝜎_’,0_, *i.e.*, the semi-principal diameter of subunit 𝑖 along the 𝑗th axis, and 𝑇 is a constant for controlling the falloff of the weight function, which is set to be 5. The weight function should be normalized if there are 𝑛 subunits in the complex. The normalized weight function can be expressed as,

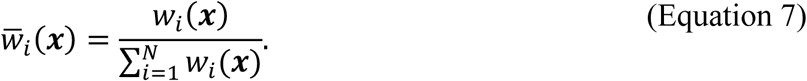

In a complex with 𝑛 subunits, the displacement of a grid point is the linear combination of movements of different subunits, which can be written as,

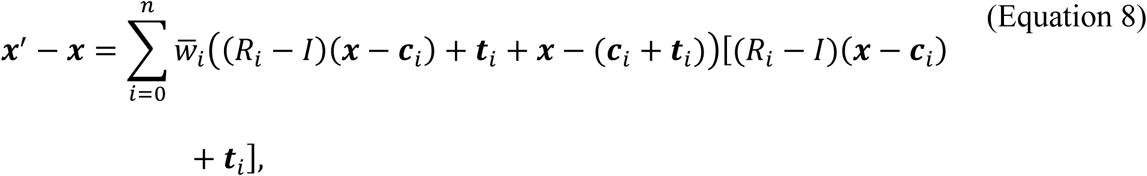

where *̄w_i_* controls the scale of the contribution of the movement of subunit 𝑖. We hence defined the deformation field 𝒈, with which the deformed grid can be represented as 𝒙’ = 𝒙 + 𝒈(𝒙).

In OPUS-ET, the rigid-body transformation parameters of subunit 𝑖, 𝑅_’_ and 𝒕_’_, are predicted by the neural network. They should be restrained to rule out unrealistic deformations. For a multi-subunit system, we assume that the translational mode of a subunit is generated by rotating the COM of subunit 𝑖 in relative to the COM of a reference subunit 𝑗, i.e., 𝒕_’_ =(𝑅*_i,t_* – 𝐼) (𝒄*_i_* – 𝒄*_j_*), where the COM 𝒄*_i_* is rotated by 𝑅*_i,t_* centering at COM 𝒄*_j_*, and the translation of subunit can be obtained as the difference between the rotated COM and the original COM. The translation defined in this way retains the distance between COMs of subunits. Users are then required to define a reference subunit for each subunit. The determination of the displacement of each subunit comes down to estimating two rotation matrices. We can further limit the range of motion by restraining the rotation matrix to be an identity matrix, which inevitably hurts the expression power of the deformation model. To maximize the capability of the deformation model under the identity restraint, the per-subunit rigid-body movement should be defined in an appropriate reference frame, *i.e.*, the 𝑧 axis should align with the rotation axis. The rotation matrix which generates the translation of COM of subunit can be further decomposed as 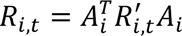, where the rotation matrix 𝐴*_i_* aligns the rotation axis of the COM of subunit to the 𝑧 axis, and 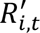 is the rotation matrix predicted by neural network. The rotation matrix of the subunit can be decomposed similarly as 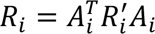_’_, where 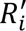 is the rotation matrix predicted by neural network. Our method adopts quaternions to represent rotation. The neural network in conformation decoder outputs the last three dimensions of the quaternion, while the first dimension of quaternion is fixed to be a positive value of 16, which can regularize it to be an identity rotation.

### Subtomogram formation model

Cryo-ET reconstructs 3D tomograms from 2D tilt-series projections whose Fourier transforms are modulated by the microscope’s contrast-transfer function (CTF). OPUS-ET models the subtomogram-formation process by combining this 3D CTF with structural deformation in the Fourier domain.

For a subtomogram 𝑋_’_, let the template volume be 𝑉, suppose 𝑋_’_ is deformed by the deformation field 𝒈 and rotated from 𝑉 by rotation matrix 𝑅_’_ with Euler angle 𝜃, the Fourier transform of a subtomogram has the following relation with respect to the Fourier transform of the template volume,

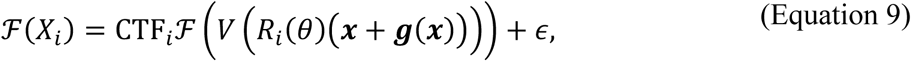

where 𝒙 is the 3D coordinate of a voxel, CTF_’_ is a 3D function in Fourier domain that describes the effects of CTF for the 2D images in the tilt series, ℱ and ℱ^./^ denote Fourier transform and inverse Fourier transform, and 𝜖 is the 3D complex Gaussian noise, which is assumed to be independent and zero-mean for each lattice point. Specifically, CTF_’_ is obtained by backprojecting the 2D CTFs of all images in the tilt series into an empty 3D volume in Fourier domain. Each 2D CTF is placed as a central slice in the 3D CTF, whose orientation is determined by its tilt angle. Reconstructing the CTF model for subtomograms by backprojection naturally handles the missing wedges as the region which is not covered by tilt series remains empty, and preserves the tilt variations of CTFs by scattering CTF from different tilts to different voxels. Another characteristic of CTF in cryo-ET is the large variations in space due to the tilting and the thickness of specimen. The tilting of the specimen holder causes the specimen swing up (overfocus) or down (underfocus) on either side of the tilt axis^13^. In addition, particles in the specimen are distributed in different heights along 𝑍 axis. Hence, subtomograms extracted from different locations of the tomogram have different defocus. Various methods have been proposed to correct the CTF gradient in cryo-ET^13^. Here, we implemented a backprojection algorithm for reconstructing 3D CTF from the per-particle CTF parameters estimated by WARP^6^. Specifically, OPUS-ET reads the defocus parameters from Self-defining Text Archiving and Retrieval (STAR) format^48^ files and reconstructs the 3DCTF for each subtomogram *ad hoc* during training.

To capture the structural heterogeneity, the reference volume in (Equation 9) is defined on a per-particle basis. We assume that the structures of subtomograms reside in a low-dimensional latent space. Since a structure can be viewed as a combination of specific composition and conformation, the latent space is assumed to be composed of a low-dimensional vector 𝒛 describing its composition and a low-dimensional vector 𝒛^!^ describing its conformation. Given a neural network 𝑉 that transforms 𝒛 into a 3D density map, and a neural network 𝒈 that transforms 𝒛^!^ into the deformation field, the Fourier transform of the subtomogram of the particle can be expressed as

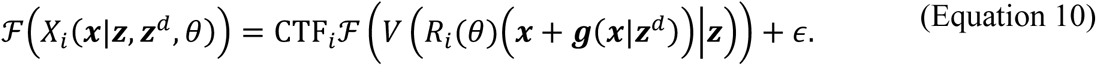

(Equation 10) defines the subtomogram formation model for the particle in cryo-ET which describes the effects of both spatial CTF variations and composition and conformation heterogeneity.

The CTF can be decomposed as a phase term, sign(CTF), which flips the sign of Fourier transform of subtomogram and leads to the spatial displacement of structure factor spacing in the image^49^, and an amplitude term, abs(CTF), which attenuates the frequency near zeros of the CTF. Hence, multiplying both sides of (Equation 10) by the phase of CTF, it can be rewritten as,

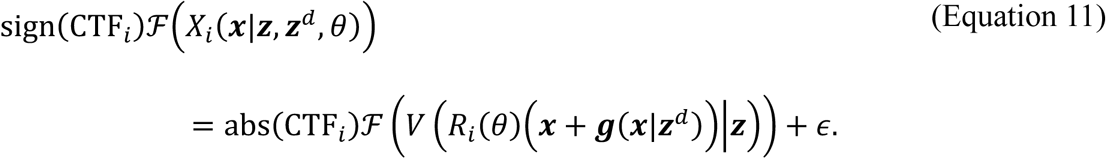

The resulting phase flipped subtomogram can be viewed as a focused subtomogram with reduced image spacing displacement, which is of higher signal to noise ratio and used for training OPUS-ET in this paper.

### Training objectives

The OPUS-ET networks are trained to minimize the difference between reconstructed and phase-flipped experimental subtomograms 𝑋_’_, that is,

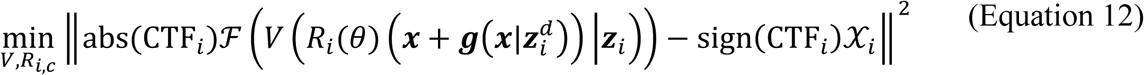

where 𝒳_’_ is the Fourier transform of 𝑋_’_, 𝒛_’_ is the composition latent code for the template volume of subtomogram 𝑖, and 𝒛^!^ is the conformation latent code for the deformation of subtomogram 𝑖. The training objective in (Equation 12) is not complete yet since the latent codes 𝒛_’_ and 𝒛^!^ of each particle remain to be inferred. We adopt the variational autoencoder (VAE) architecture where the distributions of latent codes 𝒛 and 𝒛^!^ are inferred by an encoder network 𝑓. Specifically, OPUS-ET leverages a combination of the training objective of 𝛽-VAE^37^ and the structural disentanglement prior proposed in OPUS-DSD^32^. Let the dimension of 3D volume be 𝑁^(^, the training objective of OPUS-ET is of the form,

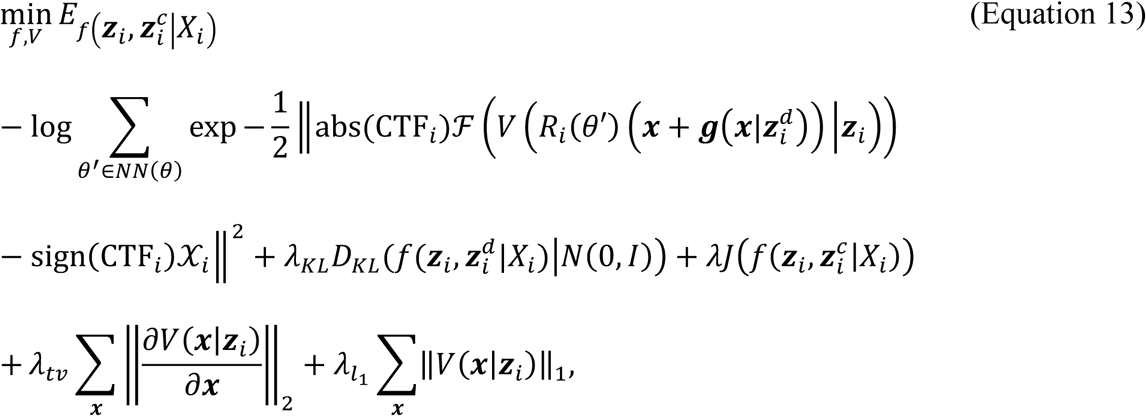

where the first term is expectation of error between the experimental subtomogram 𝑋_’_ and the reconstruction over the distributions of latent codes and the nearest neighboring poses 𝑁𝑁(𝜃) of 𝜃 in 𝑆𝑂(3), 𝑁(0, 𝐼) is the standard Gaussian distribution, 𝐷_AB_ is the KL divergence between the distributions of latent codes estimated by the encoder network 𝑓 and the standard Gaussian distribution, 𝜆_AB_ is the restraint strength for the KL divergence, 𝜆 is the restraint strength of structural disentanglement prior for the latent code of structure 𝒛, 𝜆_5C_ is the restraint strength for the total variation of structure, and 𝜆_D%_ represents the restraint strength of the 𝐿_/_norm of the structure. It is worth noting that the first expectation term in (Equation 13) is intractable, and approximated by the reparameterization trick during training ^35^.

### Experimental protocol

Tomograms for *C. reinhardtii* and *S. pombe* datasets were reconstructed in WARP^6^ using the deposited tilt-series and alignment parameters to maintain coordinate consistency with expert annotations. Candidate particles were localized by pyTOM^16^ template matching, ranked by correlation with the reference, and the highest-scoring picks retained.

3D subtomograms and per-particle 3DCTF parameters were exported from WARP and used to train OPUS-ET for purification. Poses were fixed to template-matching orientations; at this stage the conformation decoder modeled only global rigid-body motions. Clustering in the composition-latent space identified target species. The corresponding subtomograms were refined in M^4^ with image-warp correction, per-particle CTF, and pose refinement enabled. The refined particles were re-analyzed by OPUS-ET using a multi-subunit model to resolve inter-subunit conformational variation.

For benchmarking, OPUS-ET and tomoDRGN were trained on identical datasets with equal latent-space dimensionality and number of epochs; final latent representations and decoders from the last epoch were used for class selection. The same rule was followed when comparing CryoDRGN-ET.

For *S. pombe* FAS, template-matched subtomograms were refined in RELION 3.0.8^9^ using a low-pass-filtered *S. cerevisiae* FAS model. Because the complex exhibits D_3_ symmetry, symmetry expansion was applied before training. Both OPUS-ET and tomoDRGN were trained on 4,800 subtomograms with six equivalent orientations.

For *M. pneumoniae* 70S ribosomes, 22,291 subtomograms were reconstructed from the M-refined particle tilt series deposited in EMPIAR-11843^33^ using RELION 3.0.8^50^, and analyzed with OPUS-ET using the original orientations and CTF parameters. Here the conformation decoder was restricted to global rotations and translations for fair comparison.

### Training and hyperparameters’ settings

The training process of OPUS-ET followed OPUS-DSD^32^ with additional data augmentation. First, the Fourier transform of the subtomogram was blurred by a randomly sampled B-factor of [−0.25, 0.75] × 4𝜋*, the contrast and brightness were also perturbed according to the estimated SNR using random multiplicative factor in the range 1 + [−0.15,0.15] × SNR and additive factor in the range [−0.2,0.2] × SNR. The encoder generated latent codes that the decoders used to reconstruct both the 3D template and deformation field. Per-particle CTF parameters were read from STAR files to compute 3DCTFs on the fly, and reconstructed subtomograms were compared to experimental ones according to the formation model in (Equation 10).

The reconstructed subtomogram and the original subtomogram together form the reconstruction loss. We construct the training objective by combining the reconstruction loss, the priors for latent codes, and the total variation and 𝐿_/_ norms^51^ for the template volumes with specified weights as given in (Equation 13). During training, OPUS-ET uses the same batching process as in OPUS-DSD^32^ to sample subtomograms with similar projection directions as a batch.

The training losses of OPUS-ET were minimized by the Adam optimizer^52^. The learning rate of Adam was set to 3∼5 × 10^.E^. Let the SNR of dataset be SNR, the restraint strength 𝜆 for structural disentanglement prior was set to 0.5∼1 × SNR, and the restraint strength 𝜆_AB_ for KL divergence between latent codes and standard Gaussian was set to 0.5 × SNR. The SNR was estimated *ad hoc* during training by treating the reconstruction from OPUS-ET as signal and the input subtomogram with reconstruction subtracted as noise. Formally, using a batch of 𝑁 images, the SNR was estimated as,

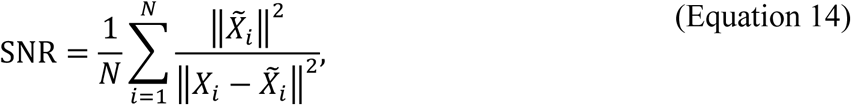

where *̃X_i_* was the subtomogram reconstructed by OPUS-ET, and *𝑋_i_* was the experimental subtomogram. ||*X_i_* – *̃X_i_*||^2^ estimated the square of the norm of noises in the experimental subtomogram, while ||*̃X_i_*||^2^ estimated the square of the norm of signals. The restraint strength 𝜆*_tv_* for the total variation of 3D reconstruction was set to 0.1, and the restraint strength 𝜆*_l_*_1_ for the 𝐿_1_ norm of 3D reconstruction was set to 0.3.

## Supporting information

Ext. Data Table 1, 2

## Data availability

We used the following publicly available datasets: EMPIAR-11830^38^ (cryo-FIB-milled lamellae of *C. reinhardtii*), EMPIAR-10988^39^ (cryo-FIB-milled lamellae of wild-type *S. pombe*), and EMPIAR-11843^33^ (Cm-treated *M. pneumoniae*). The trained weights and structural heterogeneity analysis results are available at https://doi.org/10.5281/zenodo.1263191953.

## Code availability

The source code of OPUS-ET is available at https://github.com/alncat/opusTOMO, and also at Zenodo at https://doi.org/10.5281/zenodo.1762724054.

## Acknowledgements

J.M. wants to thank the support from the National Key Research and Development Program of China (No. 2024YFA1307502), the Science and Technology Innovation Plan of Shanghai Science and Technology Commission (No. 23JS1400200), the Research Fund for International Senior Scientists (No. W2431060), and the Research Fund of Development and Reform Commission of Hunan province (No. 2507-430000-04-05-666048). The authors specially thank James Krieger for his contribution to the code improvement of OPUS-ET. The authors also thank the open-source CryoDRGN project for providing the Python framework for cryo-EM data processing upon which OPUS-ET was built.

## Author Contributions

Z.L., Q.W. and J.M. conceived the work. Z.L. designed the algorithm and implemented the software. Z.L. and X.C. performed experiments and analyses. All authors wrote and approved the final manuscript.

## Competing Interests

Q.W. is an employee of Harcam Biomedicines who is bound by confidentiality agreements that prevent the disclosure of competing interests in this work. All remaining authors declare no competing interests.

## Extended Data Figure Legends

**Ext. Data Figure 1.**
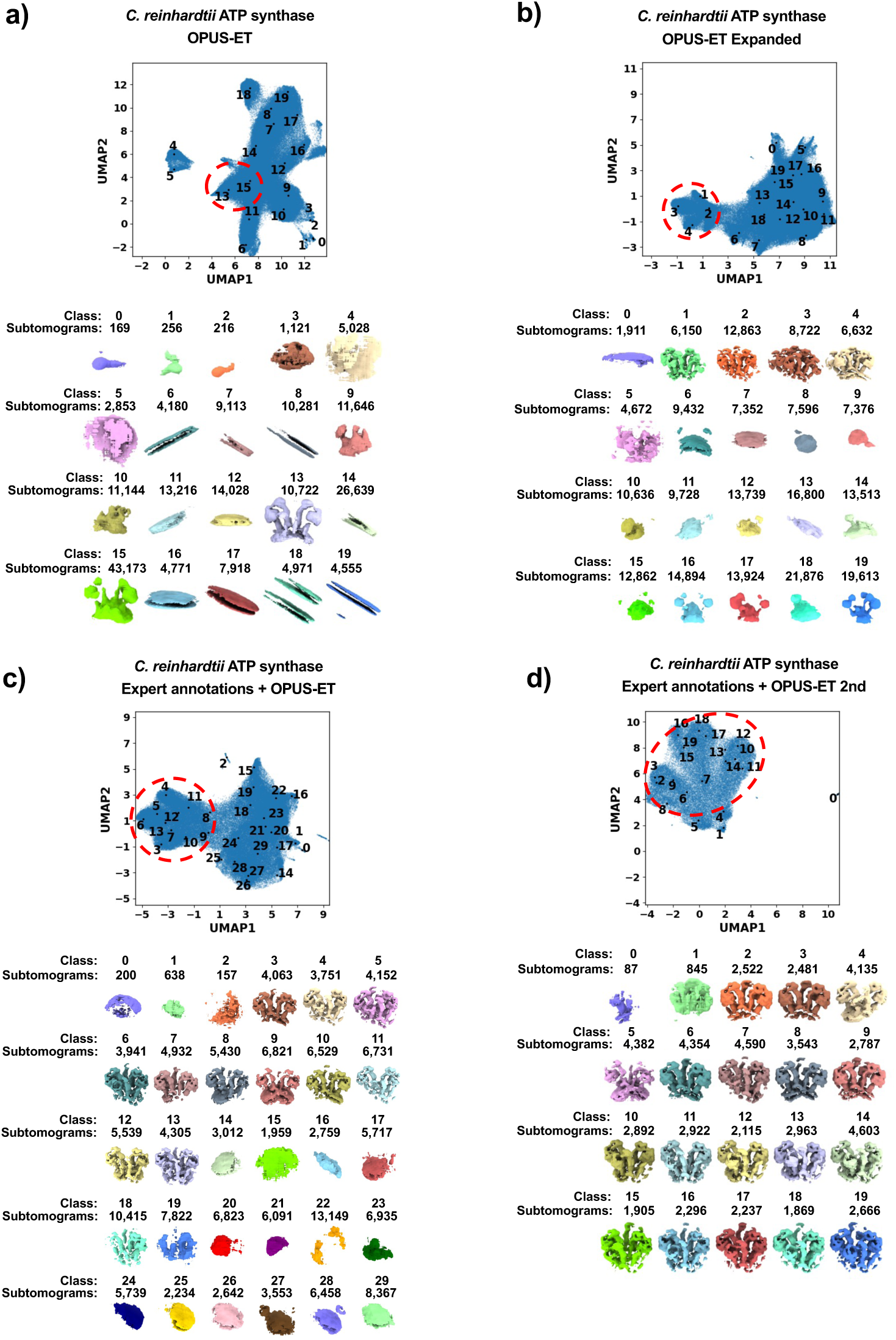
OPUS-ET’s heterogeneity analysis of *C. reinhardtii* ATP synthase dimer. **a)**. UMAP visualization of the 12-dimensional composition latent space learned by OPUS-ET on template matching result, showing the distribution of centroids for 20 classes (black dots) found by KMeans algorithm, and density maps at class centers reconstructed by the composition decoder of OPUS-ET. **b∼d**), Subsequent classification and reconstruction by OPUS-ET: **b)**. on template matching result filtered by model trained in **a)**; **c)**. on expert annotations; **d)**. on Classes 3∼13 of expert annotations in **c)**. In all panels, classes showing densities for ATP synthase are marked by red dashed ellipses.

**Ext. Data Figure 2.**
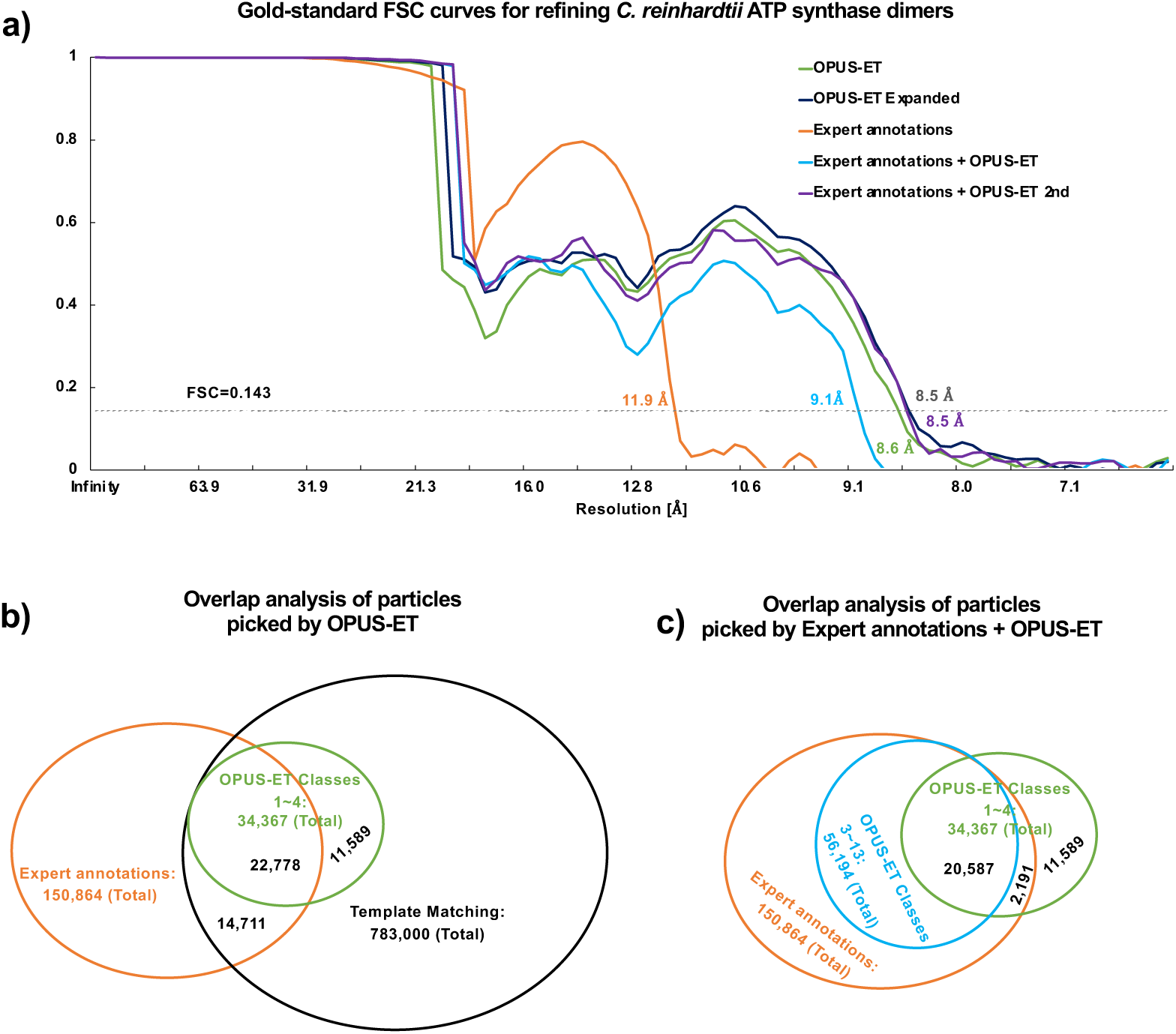
FSC curves and overlap analyses of *C. reinhardtii* ATP synthase dimer. **a)**. Gold-standard Fourier shell correlations (FSCs) curves show resolution improvement of OPUS-ET maps (8.5∼8.6 Å). The resolutions at FSC=0.143 are color labeled by datasets. **b∼c)**. Venn diagrams quantify overlap among template-matching, expert annotations, and OPUS-ET subsets. The ellipses represent 150,864 subtomograms from expert annotations (orange), 783,000 from our template matching (black), 34,367 from Classes 1∼4 in OPUS-ET’s analysis on template matching results (OPUS-ET Classes 1∼4, green), and 56,194 in Classes 3∼13 of OPUS-ET’s analysis on expert annotations (OPUS-ET Classes 3∼13, blue). The subtomograms overlapped between Expert annotations and template matching is 37,489, of which 22,778 are recovered in OPUS-ET Classes 1∼4, among which 20,587 are recovered in OPUS-ET Classes 3∼13.

**Ext. Data Figure 3.**
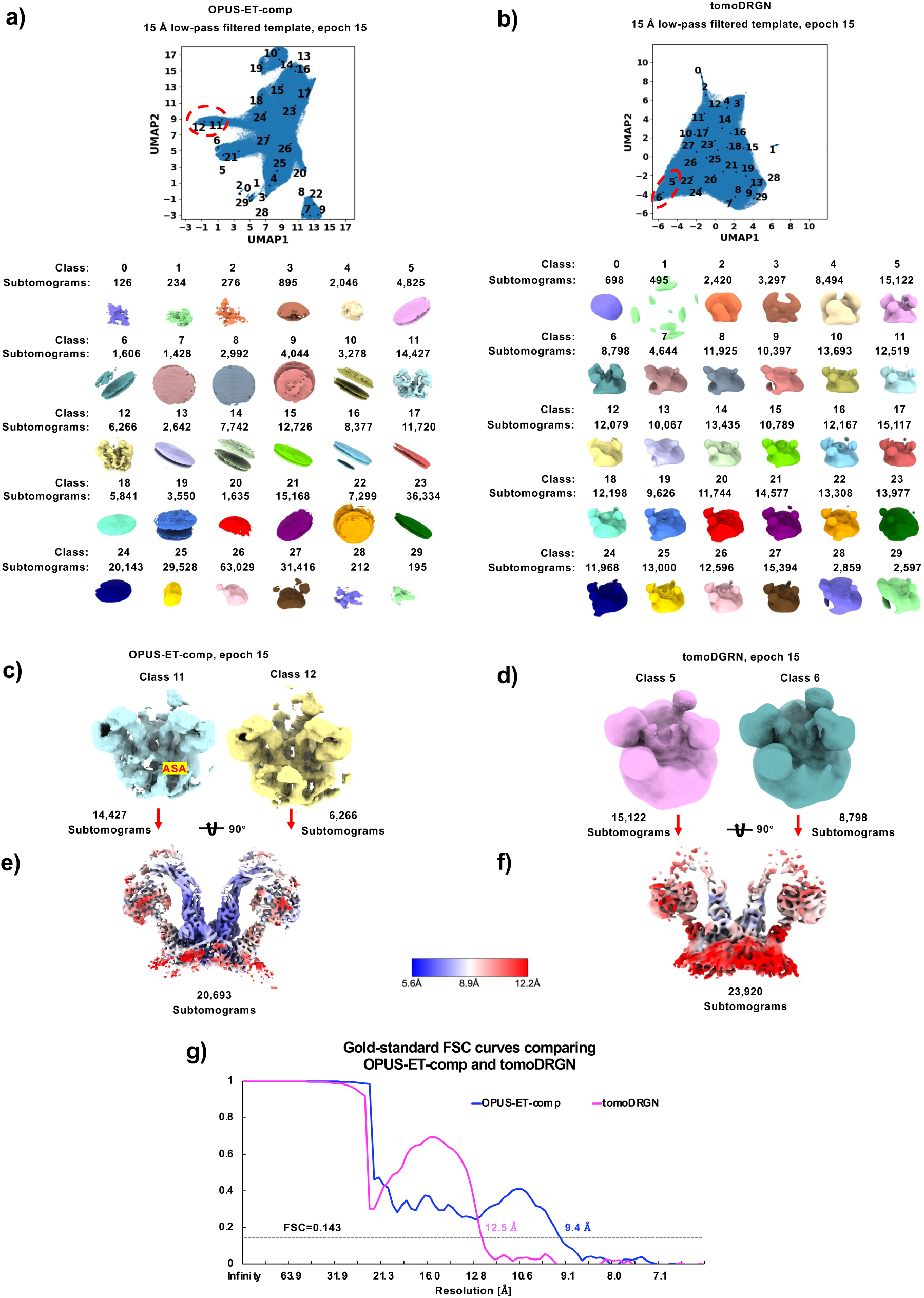
Comparison of OPUS-ET-comp and tomoDRGN on *C. reinhardtii* ATP synthase dimer. **a∼b)**. UMAP latent-space distributions after epoch 15 show clear class separation in OPUS-ET-comp **a)** but not tomoDRGN **b)**. Centroids for 30 classes are shown as black dots and classes showing densities for ATP synthase are marked by red dashed ellipses. **c∼g)**. Local-resolution maps and FSCs confirm 3.1 Å resolution advantage of OPUS-ET-comp. **c)**. Classes 11∼12 reconstructed by OPUS-ET-comp. **d)**. Classes 5∼6 reconstructed by tomoDRGN. **e∼f)**. Subtomogram average for OPUS-ET-comp Classes 11∼12 (20,693 subtomograms) and tomoDRGN Classes 5∼6 (23,920 subtomograms), respectively. The density maps are contoured at 5𝜎 and colored by local resolutions. **g)**. Gold-standard FSCs of the subtomogram averages in **e∼f**.

**Ext. Data Figure 4.**
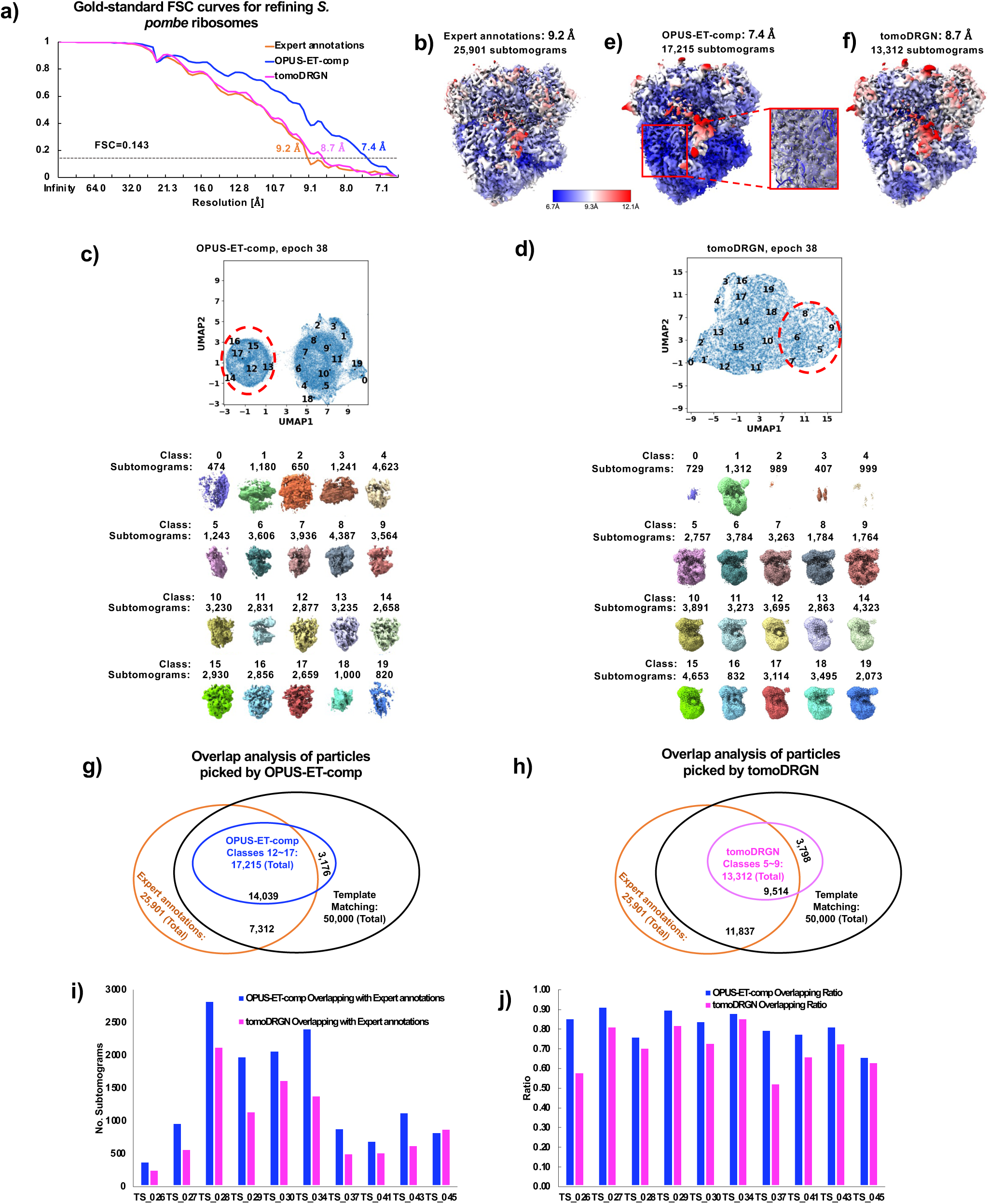
Performance of OPUS-ET on *C. reinhardtii* ATP synthase dimer with progressively low-pass filtered templates. **a∼c)**. UMAP visualizations of latent-space distributions after epoch 15 show clear class separation in template matching set with either a **a)** 15 Å, **b)** 25 Å, **c)** 35 Å low-pass filtered template. Centroids for 30 classes are shown as black dots and classes showing densities for ATP synthase are marked by red dashed ellipses. **d∼i)**. Local-resolution maps and subtomogram average show that OPUS-ET achieves sub-nanometer reconstruction of the ATP synthase dimer with templates low-pass filtered at 15 Å **(d, g)**, 25 Å **(e, h)** or 35 Å **(f, i).** The density maps are contoured at 5𝜎 and colored by local resolutions. **j)**. Gold-standard FSCs of the subtomogram averages in **g∼i**. **k)**. Summary of template-choice experiments. The table reports the template resolution, the number of selected particles, overlap with the expert annotations in **Fig.2e**, and the final reconstruction resolution.

**Ext. Data Figure 5.**
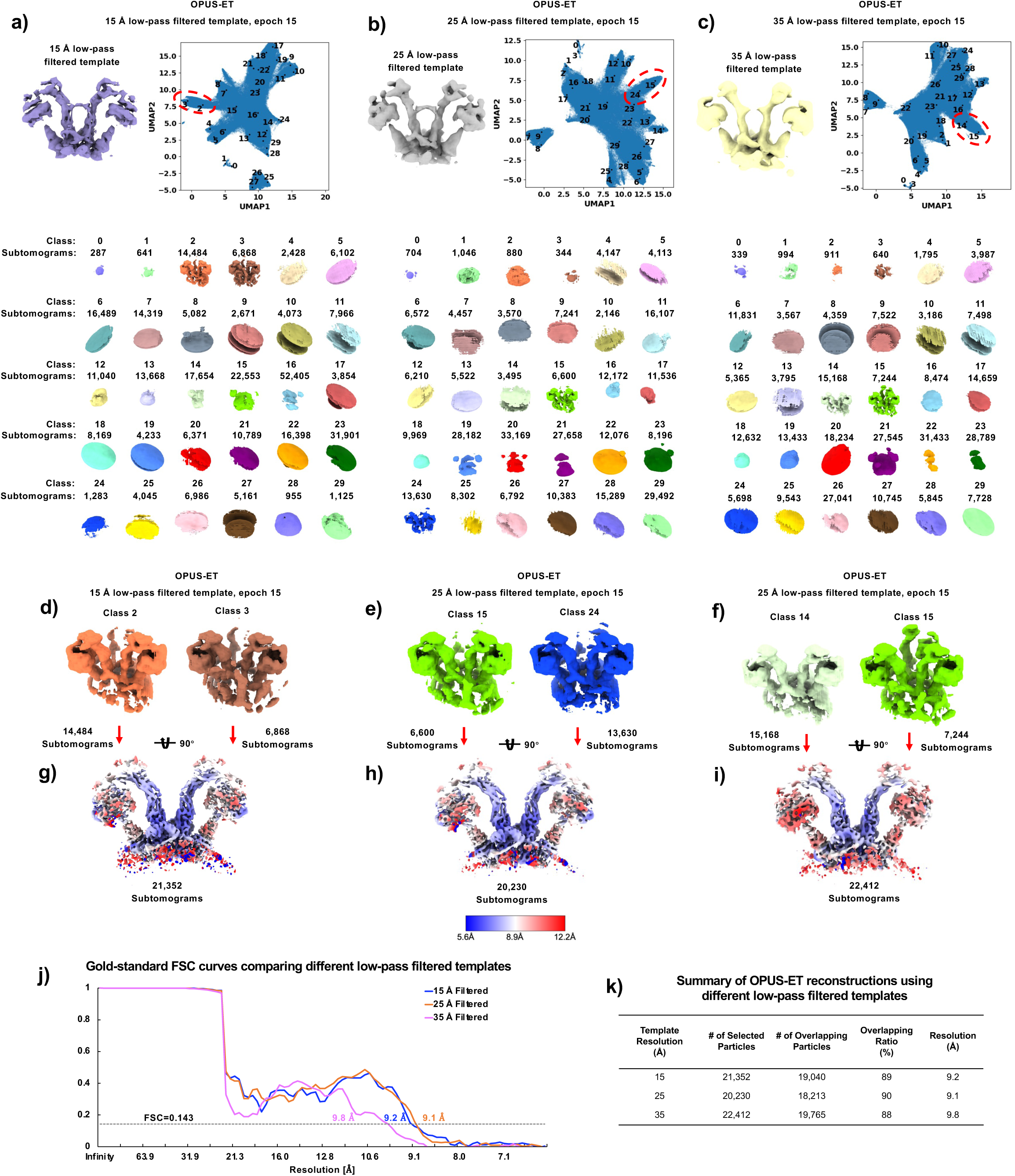
Comparison of OPUS-ET-comp and tomoDRGN on *S. pombe* 80S ribosome. **a**). Gold-standard FSCs compare expert annotations (9.2 Å), OPUS-ET-comp (7.4 Å), and tomoDRGN (8.7 Å) maps. **b)**. 9.2 Å subtomogram average of *S. pombe* 80S ribosome using 25,901 subtomograms from DeePiCt expert annotations. **c∼d)**. Composition-latent UMAPs learned by OPUS-ET-comp **c)** or tomoDRGN **d)** on template matching data at epoch 38, showing the distribution of centroids for 20 classes (black dots) and the reconstructions by the composition decoder. Red dashed ellipses highlight the classes corresponding to 80S ribosome. **e∼f)**. Local-resolution-colored maps demonstrate secondary-structure visibility in OPUS-ET-comp (red solid box zoom-in in **e**). All density maps are contoured at 4𝜎 level, and colored by local resolutions. **g∼h).** Venn diagrams quantify overlap among template-matching, expert annotations, OPUS-ET-comp **g)** and tomoDRGN **h)** subsets. The ellipses represent 25,901 subtomograms from expert annotations (orange), 50,000 from our template matching (black), 17,215 from OPUS-ET-comp Classes 12∼17 (blue), and 13,312 in tomoDRGN Classes 5∼9 (pink). The subtomograms overlapped between Expert annotations and template matching are 21,351, of which 14,039 are recovered in OPUS-ET-comp Classes 12∼17, and 9,514 are recovered in tomoDRGN Classes 5∼9. **i∼j).** Overlap subtomogram and ratio analyses across tilt-series quantify 66 % recovery of qualified ribosomes by OPUS-ET-comp vs 45 % by tomoDRGN.

**Ext. Data Figure 6.**
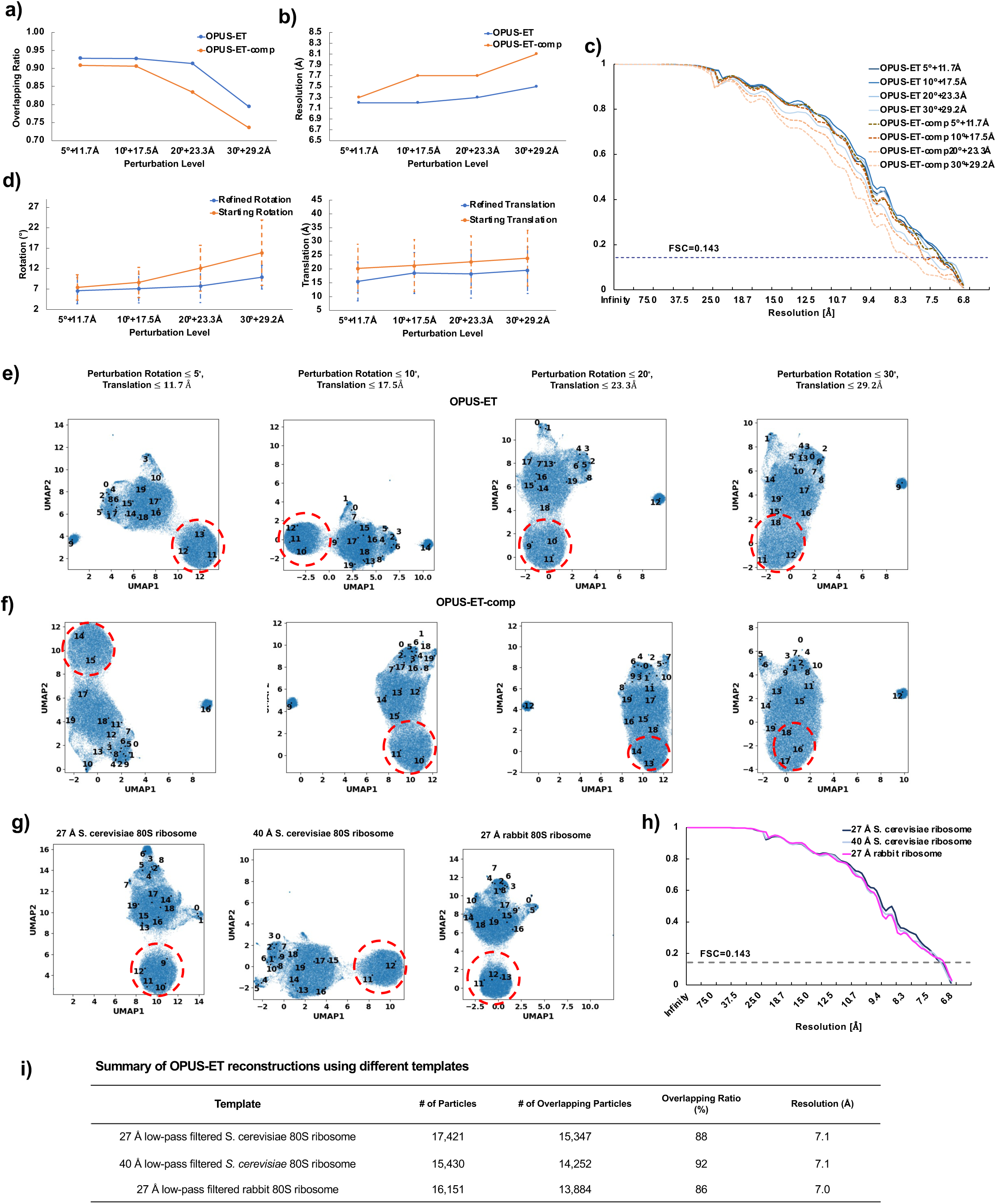
Robustness of OPUS-ET to pose perturbation and template choice in the *S. pombe* 80S ribosome dataset. **a)**. Overlap ratio between particles selected by OPUS-ET under pose perturbation and the particle set of OPUS-ET-comp in **Ext. Data Fig.5e**. **b)**. Reconstruction resolution as a function of pose perturbation level applied to the initial particle orientations and translations. Rotational perturbations (≤ 30°) and translational perturbations (≤ ∼29 Å) were introduced prior to training. Results are shown for the full OPUS-ET model and the OPUS-ET-comp configuration. **c)**. Gold-standard FSC curves corresponding to the reconstructions shown in **b)**. **d)**. Pose refinement behavior of the full OPUS-ET model under different perturbation levels. Scatter plots show starting and refined rotational and translational deviations relative to M-refined poses from **Ext. Data Fig.5e**. Refined rotation and translation are obtained by composing per-particle rotation and translation predicted by the conformation decoder with the perturbed rotation and translation. **e∼f)**. UMAP visualizations of the composition latent spaces for the pose perturbation experiments, illustrating cluster separation under increasing perturbation levels. Red dashed ellipses highlight the classes corresponding to 80S ribosome. **g)**. UMAP visualizations of the composition latent spaces for template-choice experiments. The three templates are 27 Å and 40 Å low-pass-filtered *S. cerevisiae* ribosome maps and a 27 Å low-pass-filtered rabbit ribosome map. **h)**. Gold-standard FSC curves for reconstructions obtained in **g)**. **i)**. Summary of template-choice experiments. The table reports the number of recovered particles, overlap with the OPUS-ET-comp set in **Ext. Data Fig.5e**, and the final reconstruction resolution.

**Ext. Data Figure 7.**
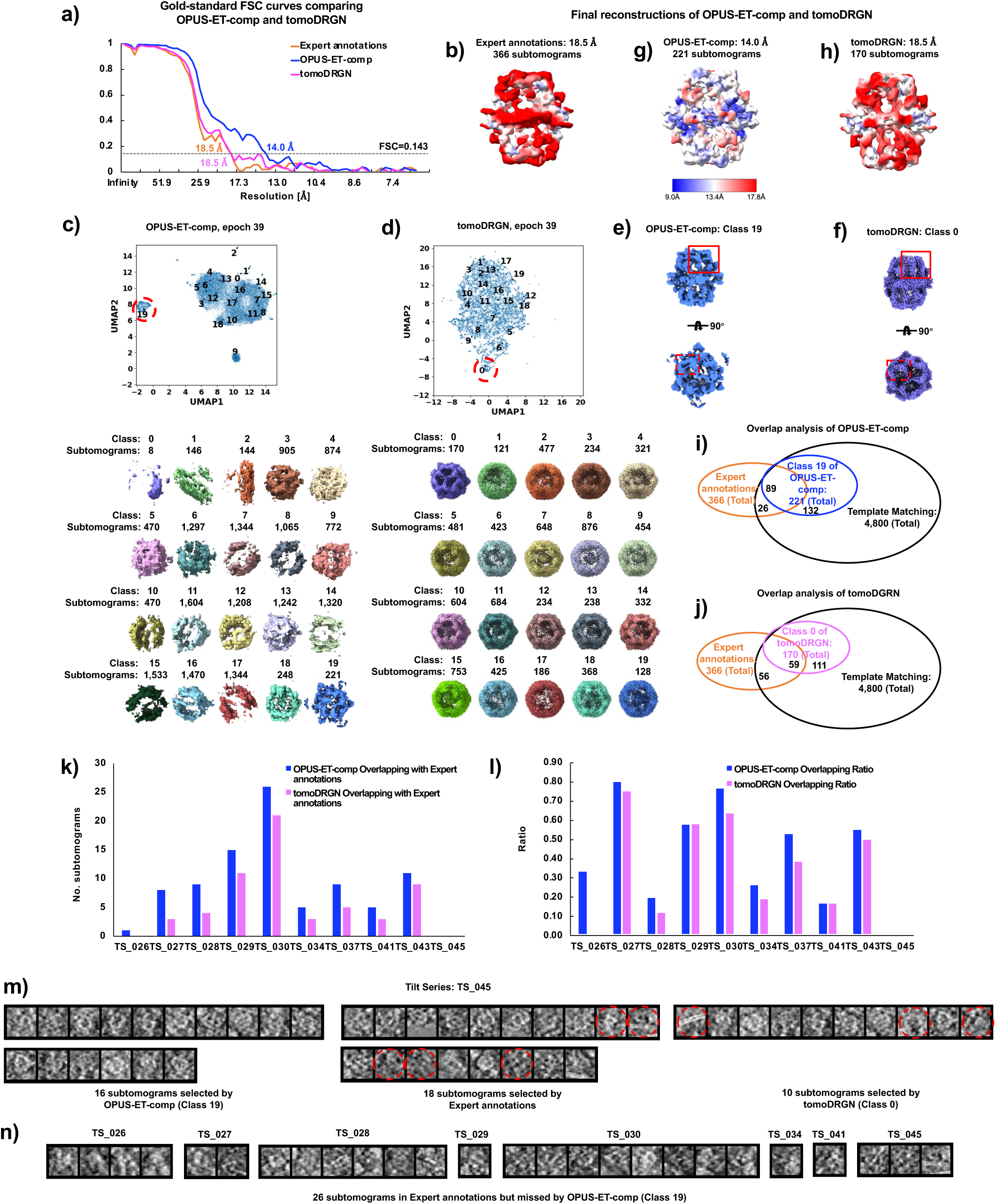
Comparison of OPUS-ET-comp and tomoDRGN on *S. pombe* FAS. **a**). Gold-standard FSCs compare expert annotations (18.5 Å), OPUS-ET-comp (14.0 Å), and tomoDRGN (18.5 Å) maps. **b)**. 18.5 Å subtomogram average of *S. pombe* FAS using 366 subtomograms from DeePiCt expert annotations. **c∼d)**. Composition-latent UMAPs learned by OPUS-ET-comp **c)** or tomoDRGN **d)** on template matching data at epoch 39, showing the distribution of centroids for 20 classes (black dots) and the reconstructions by the composition decoder. Red dashed ellipses highlight the class corresponding to FAS. **e)**. Class 19 reconstructed by OPUS-ET-comp. **f)**. Class 0 reconstructed by tomoDRGN. **g∼h)**. Local-resolution-colored maps demonstrate significantly improved map quality by OPUS-ET-comp **g)** over tomoDRGN **h)**. All density maps are contoured at 5𝜎 level, and colored by local resolutions. **i∼j).** Venn diagrams quantify overlap among template-matching, expert annotations, OPUS-ET-comp **i)** and tomoDRGN **j)** subsets. The ellipses represent 366 subtomograms from expert annotations (orange), 4,800 from our template matching (black), 221 from OPUS-ET-comp Class 19 (blue), and 170 in tomoDRGN Class 0 (pink). The subtomograms overlapped between Expert annotations and template matching are 115, of which 89 are recovered in OPUS-ET-comp Class 19, and 59 by tomoDRGN Class 0. **k∼l).** Overlap subtomogram and ratio analyses across tilt-series quantify 77 % recovery of qualified FAS particles by OPUS-ET-comp vs. 51 % by tomoDRGN. **m)**. Projections of subtomograms from the tilt series TS_045 for OPUS-ET-comp Class 19, Expert annotations and tomoDRGN Class 0. Subtomograms are CTF-corrected, 4x binned and blurred by a Gaussian (𝜎 = 1) before projecting along the 𝑍-axis. Red arrows highlight noise subtomograms. **n)**. Projections of 26 subtomograms in Expert annotations but missed by OPUS-ET-comp.

**Ext. Data Figure 8.**
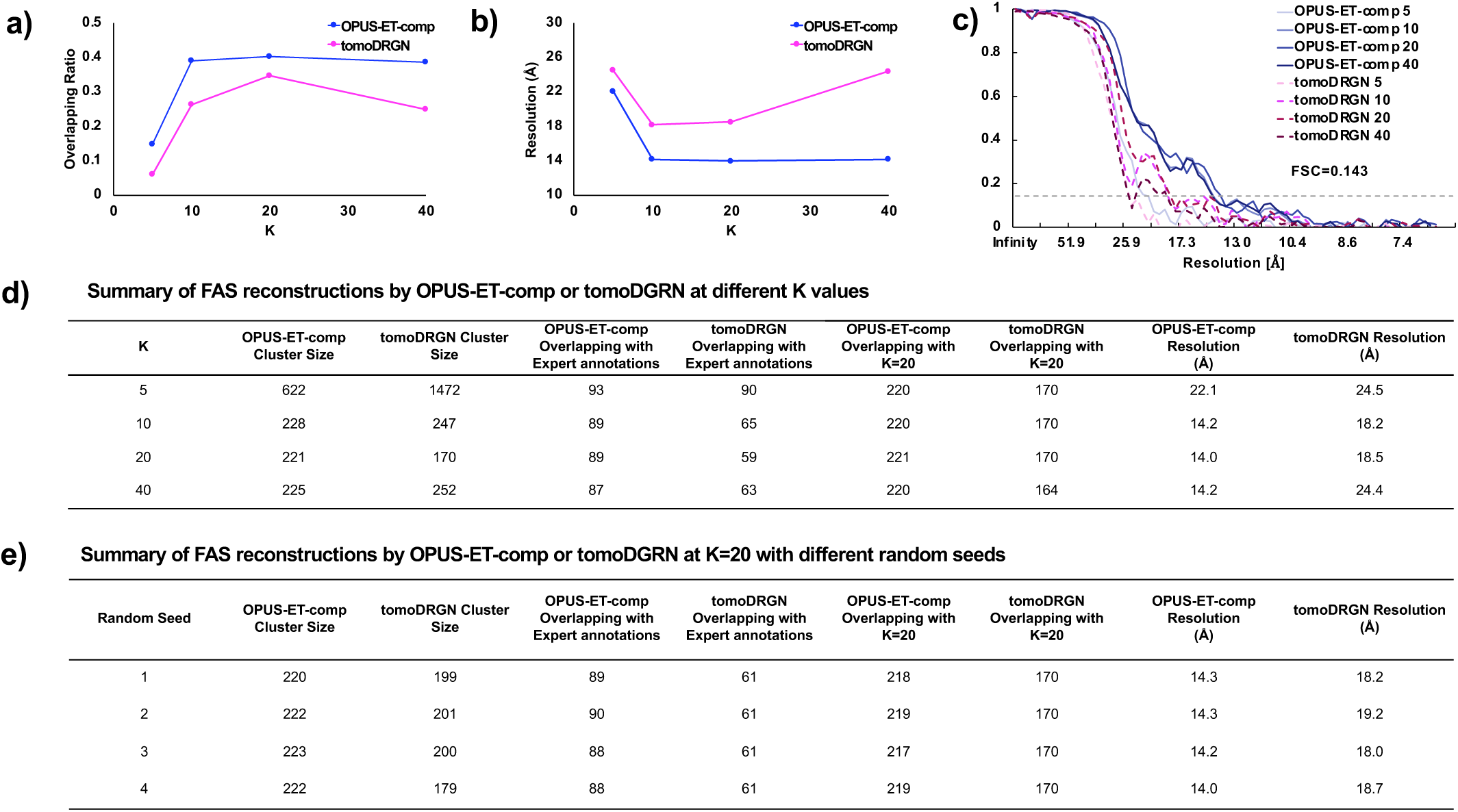
Test of OPUS-ET in recoverying FAS from the *S. pombe* dataset with a range of latent classes and random seeds. **a)**. Overlapping ratio between the selected clusters and the expert annotations as a function of the number of latent classes K, for OPUS-ET-comp and tomoDRGN. **b)**. Reconstruction resolution as a function of K for the *S. pombe* FAS dataset. Lines show the reconstruction resolution obtained after M refinement of the selected FAS clusters from OPUS-ET-comp and tomoDRGN. **c)**. Gold-standard FSC curves corresponding to the reconstructions shown in **b)**. **d∼e)**. Summary of the K-sweep analysis **d)** and random seed analysis with K=20 **e)**. Each table reports cluster size, overlap with the expert annotations, overlap with the cluster obtained at K = 20 in **Ext. Data** Fig.7, and the reconstruction resolution for OPUS-ET-comp and tomoDRGN.

**Ext. Data Figure 9.**
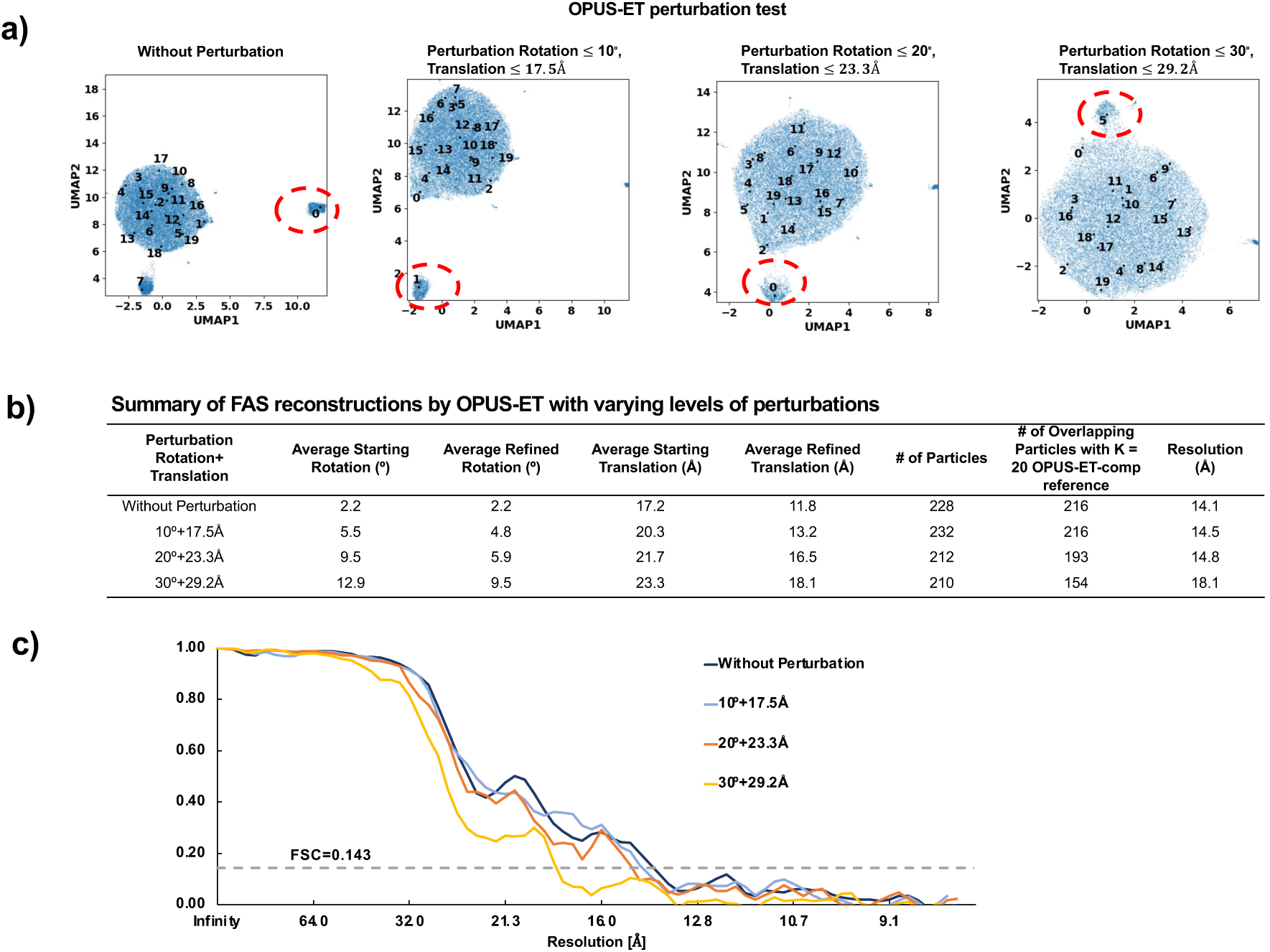
Robustness of OPUS-ET to pose perturbation in the *S. pombe* FAS dataset. **a)**. UMAP visualization of the OPUS-ET composition latent space under increasing levels of initial pose perturbation applied to FAS candidate particles. Red dashed ellipses highlight the classes corresponding to FAS. **b)**. Summary of the perturbation experiments. The table reports starting and refined rotational and translational deviations, the number of recovered particles, overlap with the OPUS-ET-comp FAS cluster in **Ext. Data Fig.7g**, and the reconstruction resolution for each perturbation level. **c)**. Gold-standard FSC curves for the reconstructions obtained under each perturbation level in **b)**.

**Ext. Data Figure 10.**
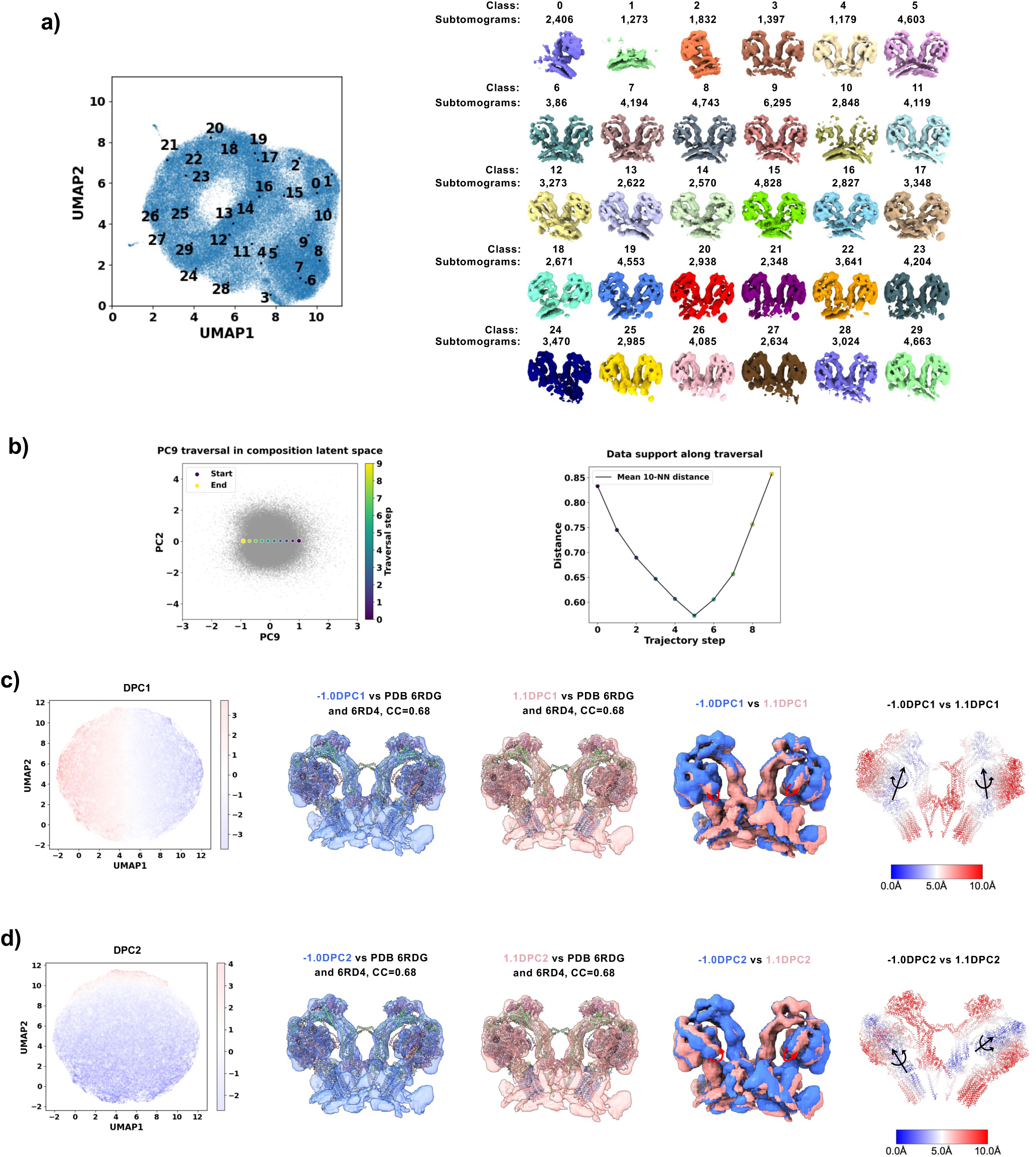
*C. reinhardtii* ATP synthase conformational modes by OPUS-ET using M-refined expanded set. **a)**. Composition-latent UMAP for expanded set (30 classes) showing the distribution of centroids for 30 classes (black dots) and their reconstructions by OPUS-ET composition decoder. **b)**. PC9 traversal shown in the latent subspace spanned by PC2 and PC9, and data support along the traversal, measured as mean distance to the 10-NN at each coordinate. **c∼d)** DPC1 and DPC2 capture bending (opposite monomer rotation) and twisting (same-direction rotation) motions with color-coded displacement maps for DPC1 **c)** and DPC2 **d). (Left)** Conformation-latent UMAP colored by their locations in DPCs (positive end = red; negative end = blue). The directions of DPC1 and DPC2 are orthogonal. **(Middle)** Density maps reconstructed by the conformation decoder of OPUS-ET using Class 5 in **a)** as the template overlaid with the individually fitted structures (PDB code 6RDG with the peripheral stalk from PDB code 6RD4). **(Right)** Superpositions of density maps at the two ends of each DPC, together with rightmost color-coded displacement maps, reveal two orthogonal inter-monomer movements. Displacement maps were calculated from per-residue displacements of the fitted structures. Black arrows indicate the rotation axes.

**Ext. Data Figure 11.**
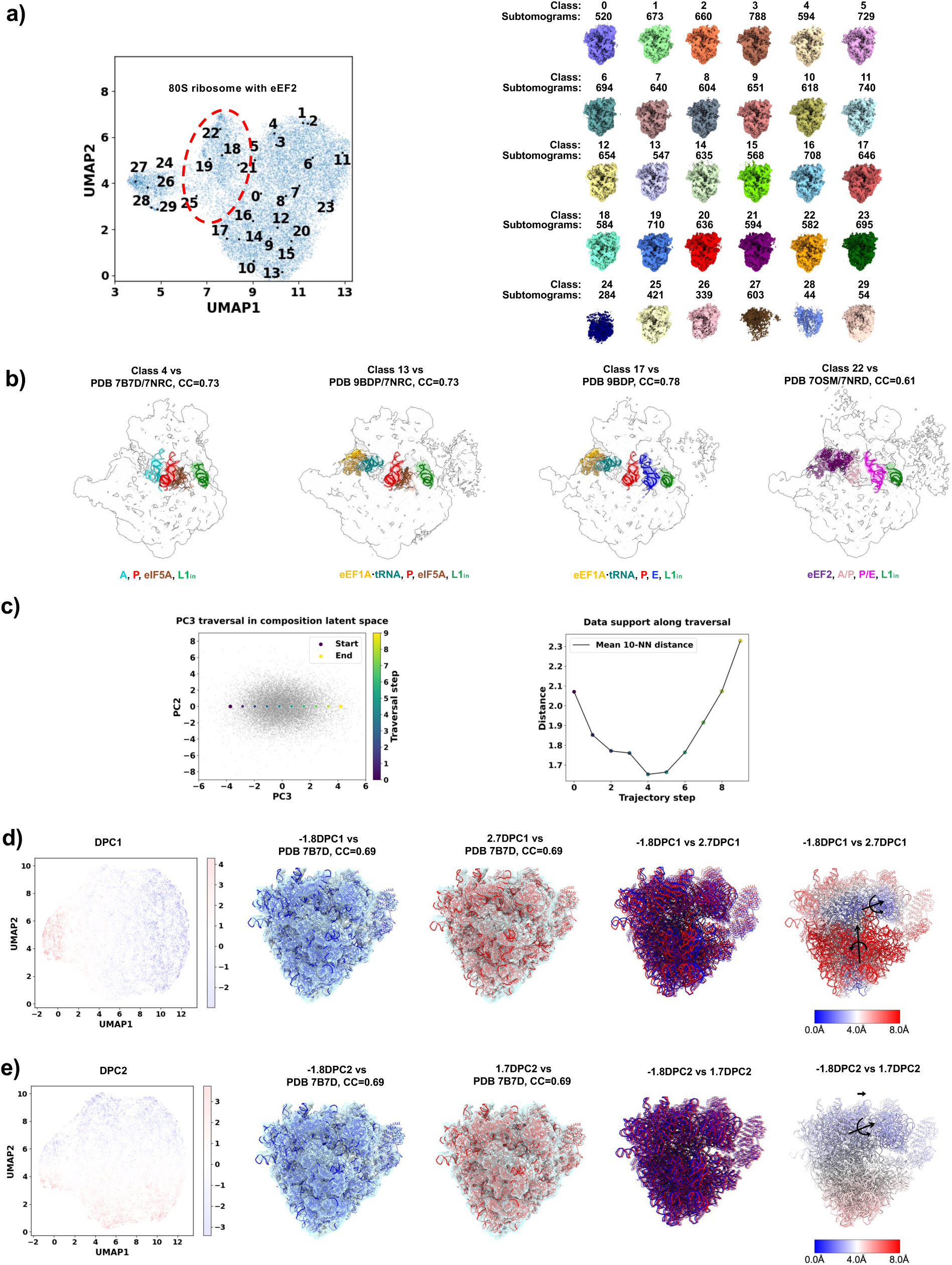
*S. pombe* 80S ribosome heterogeneity and inter-subunit dynamics. **a)**. Composition-latent UMAP of OPUS-ET on M-refined subtomogram set, showing the distribution of centroids for 30 classes (black dots) and their reconstructions by composition decoder. The classes of eEF2-containing 80S ribosome are in a red dashed ellipse. **b)** Representative tRNA translocation intermediate states revealed in **a)**. The structures of eEF1A×tRNA (orange and teal) and E-site tRNA (blue) from PDB code 9BDP, eIF5A (sienna), L1_in_ (green) and P-site tRNA (red) from PDB code 7NRC, eEF2 (indigo, PDB code 7OSM), A-site tRNA (turquoise, PDB code 7B7D), tRNAs of A/P-site (pink) and P/E-site (magenta) from PDB code 7NRD are shown as references. All densities are colored consistently with their respective models. **c)**. PC3 traversal shown in the latent subspace spanned by PC2 and PC3, and data support along the traversal, measured as mean distance to the 10-NN at each coordinate. **d∼e)** DPC1 and DPC2 capture bending (opposite monomer rotation) and twisting (same-direction rotation) motions with color-coded displacement maps for DPC1 **d)** and DPC2 **e). (Left)** Conformation-latent UMAP colored by their locations in DPCs (positive end = red; negative end = blue). The directions of DPC1 and DPC2 are orthogonal. **(Middle)** Density maps reconstructed by the conformation decoder of OPUS-ET using Class 17 in **a**) as the template overlaid with the individually fitted 40S and 60S subunits (PDB code 7B7D). **(Right)** Superpositions of density maps at the two ends of each DPC, together with rightmost color-coded displacement maps, reveal distinct inter-subunit dynamics. Displacement maps were calculated from per-residue displacements of the fitted structures. Black arrows indicate the rotation axes.

**Ext. Data Figure 12.**
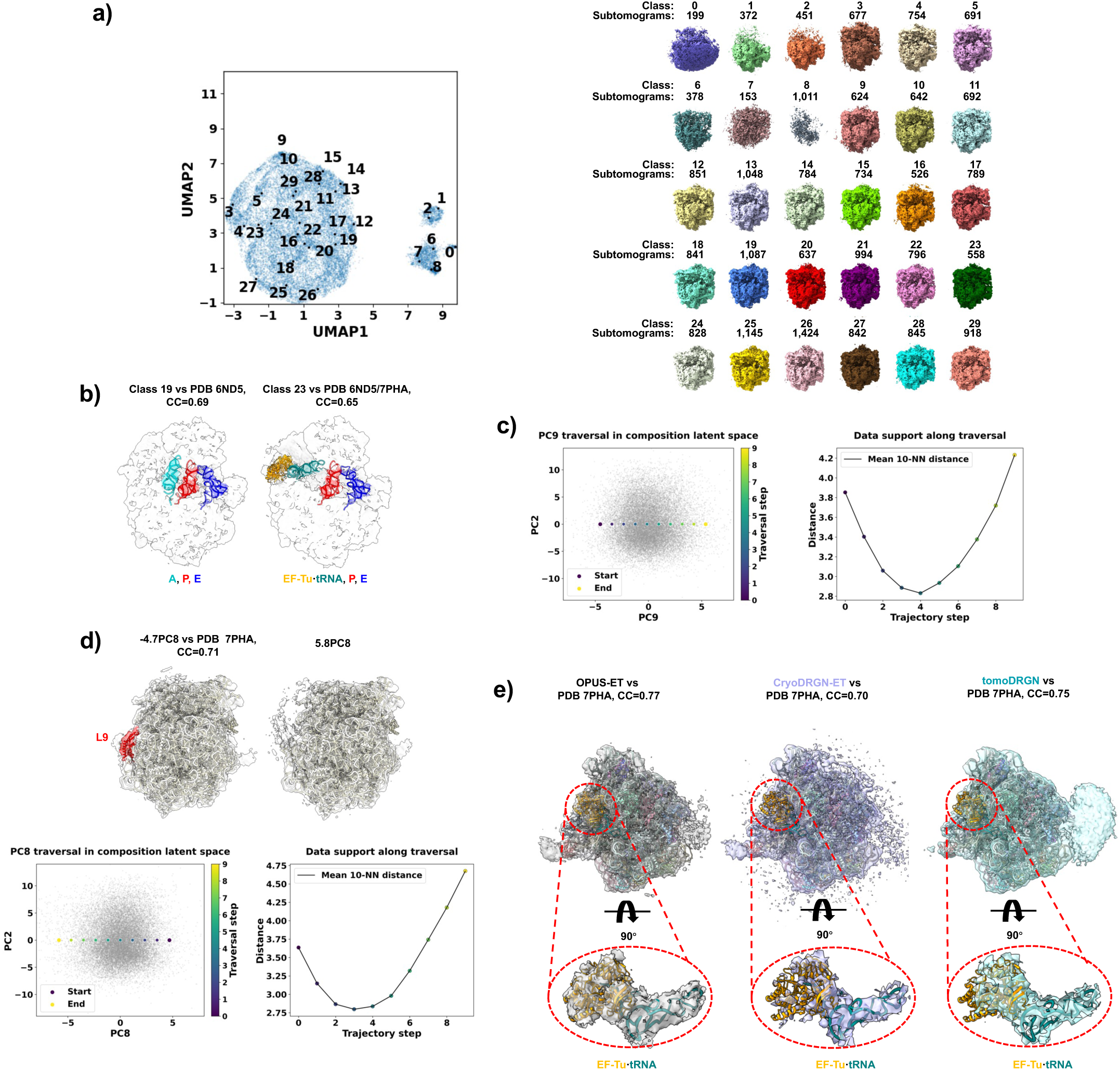
*M. pneumoniae* 70S ribosome heterogeneity. **a).** Composition-latent UMAP of OPUS-ET, showing the distribution of centroids for 30 classes (black dots) and their reconstructions by composition decoder. **b)**. Representative translocation intermediate states revealed by OPUS-ET. The structures of A- (turquoise), P- (red) and E-site (blue) tRNAs from *T. thermophilus* Cm-bound 70S ribosome (PDB code 6ND5), and EF-Tu×tRNA (orange and teal) from 70S ribosome in Cm-treated *M. pneumoniae* cell (PDB code 7PHA) are shown for reference. **c)**. PC9 traversal shown in the latent subspace spanned by PC2 and PC9, and data support along the traversal, measured as mean distance to the 10-NN at each coordinate. **d)**. (Upper) PC8 traversal reveals L9 occupancy change (red) with fitted atomic structure (PDB code 7PHA). (Lower) PC8 traversal shown in the latent subspace spanned by PC2 and PC8, and data support along the traversal, measured as mean distance to the 10-NN at each coordinate. **e**). Direct comparison shows stronger EF-Tu·tRNA densities in OPUS-ET (Left) than CryoDRGN-ET (Middle) or tomoDRGN (Right) with fitted 70S ribosome (PDB code 7PHA). Red dashed ellipses mark the densities and structures for EF-Tu×tRNA with zoom-in views. The contour levels of density maps were normalized to show similar volumes of densities for EF-Tu×tRNA in all reconstructions.

**Ext. Data Figure 13.**
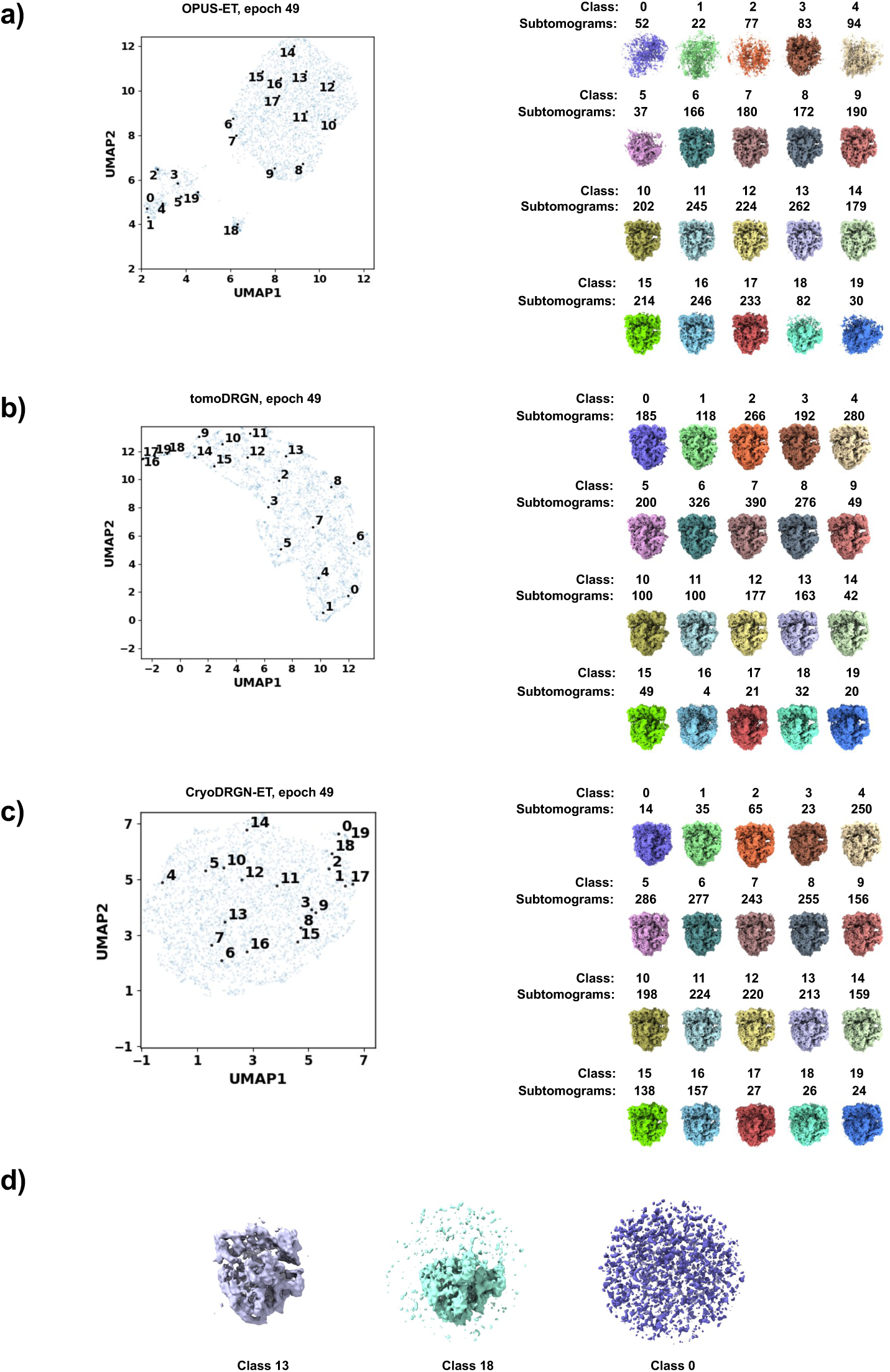
Comparison of OPUS-ET, tomoDRGN and CryoDRGN-ET on a small 70S ribosome subset. **a∼c)**. Composition-latent UMAPs on 2,990 subtomograms, showing the distribution of centroids for 20 classes (black dots) and their reconstructions by OPUS-ET **a)**, tomoDRGN **b)** and CryoDRGN-ET **c)** at epoch 49. **d)**. Subtomogram averages for three representative classes by OPUS-ET in **a)**. The subtomogram averages are blurred by Gaussian kernel (𝜎 = 2).

## Supplementary Video Legends

**Suppl. Video 1. ATP synthase PC9 rotation.** Traversal of PC9 reveals concerted 120° γ-subunit rotation with ∼30° β-head counter-motion from Substate 3C to 1A in *C. reinhardtii* ATP synthase. γ subunit of Substate 1A is shown to mark the locations of densities in Substate 1A.

**Suppl. Video 2. ATP synthase PC8 rotation.** Traversal of PC8 in another training run reveals similar counter rotations as in **Suppl. Video 1** for *C. reinhardtii* ATP synthase. γ subunit of Substate 1A is shown to mark the locations of densities in Substate 1A.

**Suppl. Video 3. ATP synthase DPC1 bending.** Traversal of DPC1 reveals the bend of ATP synthase dimer from opposite monomer rotations along parallel axes.

**Suppl. Video 4. ATP synthase DPC2 twisting.** Traversal of DPC2 reveals the twist of ATP synthase dimer from monomers rotating in same direction about orthogonal axes.

**Suppl. Video 5. 80S ribosome PC3 tRNA translocation.** Traversal of PC3 reveals the continuous transition from A/T–P to A/P–P/E states with L1-stalk closure and 40S counter-clockwise rotation. The atomic models for eEF1A×tRNA (orange and teal), A-site tRNA (green), P-site tRNA (red) and E-site tRNA (blue) are shown for reference.

**Suppl. Video 6. 80S ribosome PC4 tRNA translocation.** Traversal of PC4 in another training run reveals the continuous transition from A/T–P to A/P–P/E states with L1-stalk closure and 40S counter-clockwise rotation. The atomic models for eEF1A×tRNA (orange and teal), A-site tRNA (green), P-site tRNA (red) and E-site tRNA (blue) are shown for reference.

**Suppl. Video 7. 80S ribosome DPC1 counter-clockwise rotation.** Traversal of DPC1 reveals that 40S and 60S subunits rotate in opposite directions around orthogonal axes.

**Suppl. Video 8. 80S ribosome DPC2 co-rotation.** Traversal of DPC2 reveals coordinated translation and rotation of 40S and 60S subunits along a shared axis.

**Suppl. Video 9. 70S ribosome PC9 translation continuum revealed by OPUS-ET.** Traversal of PC9 reveals tRNA delivery and E-site release with L1-stalk movement (L1_in_ → L1_out_). The atomic models for EF-Tu×tRNA (orange and teal), A-site tRNA (turquoise), P-site tRNA (red) and E-site tRNA (blue) are shown for reference.

**Suppl. Video 10. Comparison of CryoDRGN-ET and tomoDRGN translation trajectories.** The dynamics reconstructed by CryoDRGN-ET (Left) used the first 32 deposited volumes. The dynamics reconstructed by tomoDRGN (Right) used deposited volumes for Classes 23∼32 from the 100 classes clustered by KMeans algorithm. The atomic models for EF-Tu×tRNA (orange and teal), A-site tRNA (turquoise), and P-site tRNA (red) are shown for reference.

## Notes

### Competing Interest Statement

Q.W. is an employee of Harcam Biomedicines who is bound by confidentiality agreements that prevents the disclosure of competing interests in this work. All remaining authors declare no competing interest.

### Summary of Updates

added new experiments using alternative templates and different network configurations; author affiliations updated; Supplemental files updated.

