## Supplementary material for "OPUS-ET: Resolving Compositional and Conformational Heterogeneities of Biomolecules in Cryo-Electron Tomography": Ext. Data Table 1, 2

### Ext. Data Table 1. Benchmark performance of OPUS-ET and reference methods across four cellular systems

| **Dataset/ EMPIAR** | **No. Tomograms** | **Method** | **Selected subtomograms** | **Final resolution (Å)^1^** | **Resolution improvement (Å)** | **Runtime (minutes) / epoch^2^** | **Key findings** |
| --- | --- | --- | --- | --- | --- | --- | --- |
| *C. reinhardtii* ATP synthase dimer,  EMPIAR- 11830 | 261 | OPUS-ET | 46,745 | 8.5 | +3.4 over expert | ~312 | Direct enrichment from noisy template-matching; sub-nanometer map resolving F₀–F₁ coupling and ASA architecture; discovered normal-mode-like bending and twisting of dimers |
|  |  | Expert annotations | 150,864 | 11.9 | – | NA | Labor-intensive multi-round 3D classification |
|  | 100 | OPUS-ET | 20,693 | 9.4 | +3.1 over tomoDRGN | ~120 | Resolved peripheral-stalk and ASA subunits |
|  |  | tomoDRGN | 23,920 | 12.5 | – | ~276 | Poor separation of oligomeric states; low overall quality of reconstruction |
| *S. pombe*  80S ribosome, EMPIAR- 10988 | 10 | OPUS-ET | 17,215 | 7.4 | +1.8 over expert;  +1.3 over tomoDRGN | ~22 | Recovered 66% of qualified ribosomes; captured eEF1A/eEF2 intermediates; resolved secondary-structure elements |
|  |  | Expert annotations | 25,901 | 9.2 | – | NA | Manual expert curation; low local resolution |
|  |  | tomoDRGN | 13,312 | 8.7 | +0.5 over expert | ~45 | Less discriminative latent space; blurred RNA features; recovered 45% of qualified ribosomes |
| *S. pombe* FAS, EMPIAR-10988 | 10 | OPUS-ET | 221 | 14.0 | +4.5 over expert and tomoDRGN | ~12 | Recovered low-abundance FAS (77%); improved local resolution |
|  |  | Expert annotations | 366 | 18.5 | – | NA | Low local resolution; incomplete densities |
|  |  | tomoDRGN | 170 | 18.5 | +0 over expert | ~16 | Incomplete map; misclassified membrane fragments; 51% recovery of qualified subtomograms |
| *M. pneumoniae* 70S ribosome, EMPIAR-11843 | 65 | OPUS-ET | 22,291 | 3.5 | – | ~5.53 | Reconstructed a more complete translation trajectory automatically; revealed concerted conformational changes during transitions; identified new translocation-intermediate states |
|  |  | CryoDRGN-ET / tomoDRGN | 18,466/22,291 | 3.6/3.5 | – | NA | Required manual interpolation or KMeans sampling to visualize motions |
|  | 8 | OPUS-ET | 2,990 | – | – | ~0.90 | Discriminated complete 70S from 50S and noise |
|  |  | CryoDRGN-ET / tomoDRGN | 2,990 | – | – | ~0.46 / ~1.50 | Less discriminative latent space |

^1^All resolutions were determined using the gold-standard FSC = 0.143 criterion. ^2^Runtimes (in minutes) represent average wall-clock duration on four NVIDIA V100 GPUs with 128³-voxel inputs. “NA” denotes “Not applicable” because the relevant information was extracted from prior publications.

### Ext. Data Table 2. Training commands and peak memory of OPUS-ET and reference methods across four cellular systems

| **Dataset/ EMPIAR** | **Method** | **Training commands^1^** | **Peak Memory** |
| --- | --- | --- | --- |
| *C. reinhardtii* ATP synthase dimer,  EMPIAR- 11830 | OPUS-ET on template matching results | torchrun --nproc_per_node=4 train_tomo_dist atps3d.star --poses atps3d_pose_euler.pkl -n 20 -b 16 --zdim 12 --lr 3.0e-5 --num-gpus 4 --multigpu --beta-control 0.5 -o . -r ../../atp/mask_ref.mrc --split deep.pkl --lamb 0.5 --bfactor 3. --valfrac 0.1 --templateres 144 --tmp-prefix tmp --datadir /work/jpma/luo/tomo/visual_pro/metadata/warp_tiltseries/ --angpix 3.92 --downfrac 1. --warp --tilt-range 69 --tilt-step 3 --ctfalpha 0.5 --ctfbeta 0.5 --estpose | 22GB per GPU |
|  | OPUS-ET on high resolution subtomogram averaging result | torchrun --nproc_per_node=4 -m cryodrgn.commands.train_tomo_dist matpfull_expanded.star --poses matpfull_expanded_pose_euler.pkl -n 50 -b 8 --zdim 12 --lr 3.5e-5 --num-gpus 4 --multigpu --beta-control 0.5 -o . -r ../atpmask.mrc --split deep_splt.pkl --lamb 0.5 --bfactor 10. --valfrac 0.1 --templateres 192 --tmp-prefix tmp --datadir /work/jpma/luo/tomo/visual_pro/metadata/warp_tiltseries/ --angpix 2.72 --downfrac 1. --warp --tilt-range 69 --tilt-step 3 --estpose --ctfalpha 0.5 --ctfbeta 0.5 --masks ../mask2new.pkl --accum-step 2 | 28GB per GPU |
|  | OPUS-ET on template matching results of 100 tomograms | torchrun --nproc_per_node=4 train_tomo_dist atps1003d.star --poses atps1003d_pose_euler.pkl -n 16 -b 10 --zdim 8 --lr 3.0e-5 --num-gpus 4 --multigpu --beta-control 0.5 -o . -r ../../atp/mask_ref.mrc --split deep.pkl --lamb 0.5 --bfactor 3. --valfrac 0.05 --templateres 128 --tmp-prefix tmp --datadir /work/jpma/luo/tomo/visual_pro/metadata/warp_tiltseries/ --angpix 3.92 --downfrac 1. --plot --warp --tilt-range 69 --tilt-step 3 --ctfalpha 0.5 --ctfbeta 0.5 | 10GB per GPU |
|  | tomoDRGN on template matching results of 100 tomograms | tomodrgn train_vae ../../warp_tiltseries/atps1402d_optimisation_set.star --outdir zatp100/ --zdim 8 --uninvert-data --num-epochs 16 --l-dose-mask --recon-dose-weight --recon-tilt-weight --batch-size 10 --lazy --num-workers 2 --prefetch-factor 2 --persistent-workers --multigpu --source-software warptools | 7GB per GPU |
| *S. pombe*  80S ribosome, EMPIAR- 10988 | OPUS-ET on template matching results | dsd train_tomo ribotm.star --poses ribotm_pose_euler.pkl -n 40 -b 10 --zdim 12 --lr 3.e-5 --num-gpus 4 --multigpu --beta-control 0.5 -o . -r ../zribotm_pose/mask.mrc --split deep.pkl --lamb 0.5 --bfactor 3. --valfrac 0.05 --templateres 128 --tmp-prefix tmp --datadir /work/jpma/luo/tomo/warp_DEF/metadata/warp_tiltseries/ --angpix 3.37 --downfrac 1. --warp --tilt-range 50 --tilt-step 2 --ctfalpha 0. --ctfbeta 1. | 10GB per GPU |
|  | tomoDRGN on template matching results | tomodrgn train_vae ../../warp_tiltseries/ribotm2d_optimisation_set.star --outdir z12/ --zdim 12 --uninvert-data --num-epochs 40 --l-dose-mask --recon-dose-weight --recon-tilt-weight --batch-size 10 --lazy --num-workers 2 --prefetch-factor 2 --persistent-workers | 7GB per GPU |
|  | OPUS-ET on pose perturbations and template dependence of template matching result with full model | dsd train_tomo ribotm.star --poses ribotm_pose_euler.pkl -n 40 -b 12 --zdim 12 --lr 3.5e-5 --num-gpus 4 --multigpu --beta-control 0.5 -o . -r ../zribotm_pose/mask.mrc --split deep.pkl --lamb 0.5 --bfactor 3. --valfrac 0.05 --templateres 128 --tmp-prefix tmp --datadir /work/jpma/luo/tomo/warp_DEF/metadata/warp_tiltseries/ --angpix 3.37 --downfrac 1. --warp --tilt-range 50 --tilt-step 2 --ctfalpha 0. --ctfbeta 1. --estpose --accum-step 2 | 12GB per GPU |
|  | OPUS-ET on pose perturbations of template matching result with composition-only model | dsd train_tomo ribotm.star --poses ribotm_pose_euler.pkl -n 40 -b 12 --zdim 12 --lr 3.5e-5 --num-gpus 4 --multigpu --beta-control 0.5 -o . -r ../zribotm_pose/mask.mrc --split deep.pkl --lamb 0.5 --bfactor 3. --valfrac 0.05 --templateres 128 --tmp-prefix tmp --datadir /work/jpma/luo/tomo/warp_DEF/metadata/warp_tiltseries/ --angpix 3.37 --downfrac 1. --warp --tilt-range 50 --tilt-step 2 --ctfalpha 0. --ctfbeta 1. --accum-step 2 | 12GB per GPU |
|  | OPUS-ET on high resolution subtomogram averaging result | torchrun --nproc_per_node=4 -m cryodrgn.commands.train_tomo_dist ../zribo_test/matching80s.star --poses ../zribo_test/matching80s_pose_euler.pkl -n 60 -b 8 --zdim 12 --lr 4e-5 --num-gpus 4 --multigpu --beta-control 0.5 -o . -r ../zribo_test/mask.mrc --split deep.pkl --lamb 1.0 --bfactor 4. --valfrac 0.05 --templateres 160 --tmp-prefix tmp --datadir /work/jpma/luo/tomo/warp_DEF/metadata/warp_tiltseries/ --angpix 3.37 --downfrac 1. --warp --tilt-range 50 --tilt-step 2 --ctfalpha 0. --ctfbeta 1. --accum-step 1 --estpose --masks ../mask_params.pkl --encoderres 13 | 16GB per GPU |
| *S. pombe* FAS, EMPIAR-10988 | OPUS-ET on template matching result | dsd train_tomo ../zfastm/fastmexpanded.star --poses ../zfastm/fastmexpanded_pose_euler.pkl -n 40 -b 10 --zdim 10 --lr 3.e-5 --num-gpus 4 --multigpu --beta-control 0.5 -o . -r ../zribotmt/mask.mrc --split deep.pkl --lamb 0.5 --bfactor 3. --valfrac 0.05 --templateres 128 --tmp-prefix tmp --datadir /work/jpma/luo/tomo/warp_DEF/metadata/warp_tiltseries/ --angpix 3.37 --downfrac 1. --warp --tilt-range 50 --tilt-step 2 --ctfalpha 0. --ctfbeta 1. | 10GB per GPU |
|  | tomoDRGN on template matching result | tomodrgn train_vae ../../warp_tiltseries/fastm2d_optimisation_set.star --outdir zfas/ --zdim 10 --uninvert-data --num-epochs 40 --l-dose-mask --recon-dose-weight --recon-tilt-weight --batch-size 10 --lazy --num-workers 2 --prefetch-factor 2 --persistent-workers --source-software warptools --multigpu | 7GB per GPU |
|  | OPUS-ET on pose perturbations of template matching result with full model | torchrun --nproc_per_node=4 cryodrgn.commands.train_tomo_dist fasexpanded.star --poses fasexpanded_pose_euler.pkl -n 40 -b 10 --zdim 10 --lr 3.0e-5 --num-gpus 4 --multigpu --beta-control 0.5 -o . -r ../zribotmt/mask.mrc --split deep.pkl --lamb 0.5 --bfactor 3. --valfrac 0.05 --templateres 128 --tmp-prefix tmp --datadir /work/jpma/luo/tomo/warp_DEF/metadata/warp_tiltseries/ --angpix 3.37 --downfrac 1. --warp --tilt-range 50 --tilt-step 2 --ctfalpha 0. --ctfbeta 1. --estpose --accum-step 2 | 10GB per GPU |
| *M. pneumoniae* 70S ribosome, EMPIAR-11843 | OPUS-ET on full dataset | dsd train_tomo ../op64/subtomos.star --poses ../op64/subtomos_pose_euler.pkl -n 40 -b 8 --zdim 12 --lr 4e-5 --num-gpus 4 --multigpu --beta-control 0.5 -o . -r ../opus/mask.mrc --split ribola_split.pkl --lamb 0.5 --bfactor 2. --valfrac 0.1 --templateres 192 --datadir /work/jpma/luo/tomo/10499/drgn/subtomos/ --angpix 1.7 --downfrac 0.9 --estpose | 28GB per GPU |
|  | CryoDRGN-ET on 8 tomograms | cryodrgn train_vae test8.star --ctf test8etctf.pkl --poses test8etpose.pkl --encode-mode tilt --dose-per-tilt 3.0 --zdim 8 -n 50 -o drgnet8/ --multigpu | 7GB per GPU |
|  | OPUS-ET on 8 tomograms | torchrun --nproc_per_node=4 -m cryodrgn.commands.train_tomo_dist train_tomo ../op8/subtomos.star --poses ../op8/subtomos_pose_euler.pkl -n 50 -b 8 --zdim 8 --lr 5e-5 --num-gpus 4 --multigpu --beta-control 0.5 -o . -r ../opus/mask.mrc --split ribola_split.pkl --lamb 1. --bfactor 2. --valfrac 0.0 --templateres 176 --datadir /work/jpma/luo/tomo/10499/drgn/subtomos96/subtomos/ --angpix 3.7 --downfrac 1. --estpose --ctfalpha 0 --ctfbeta 1 | 20GB per GPU |
|  | tomoDRGN on 8 tomograms | tomodrgn train_vae test8.star --zdim 8 --outdir drgn8 --num-epochs 50 --multigpu | 4GB per GPU |

^1^--templateres specifies the output size of convolutional neural network (in voxels) used for training. --angpix is the pixel size in Ångströms. -n and -b specify the number of training epochs and batch size per GPU, respectively. --zdim controls the dimensionality of the composition latent space. --lr is the learning rate. --lamb controls the weight of the structural disentanglement prior. --bfactor applies a B-factor blurring in CTF to the decoder reconstruction during training to adjust the frequency weighting of the reconstruction loss. --valfrac sets the fraction of particles held out for validation. --ctfalpha controls the degree of CTF correction applied to the phase-flipped experimental subtomogram, scaling the Fourier transform by |CTF|^alpha^. The default value of 0 is equivalent to phase flipping only; higher values apply progressively stronger amplitude correction. --ctfbeta controls the degree of CTF correction applied to the decoder reconstruction, scaling it by |CTF|^beta^. The default value of 1 applies full CTF correction. --estpose enables pose correction by the conformation decoder during training. --masks specifies the rigid-body subunit partition file for the conformation decoder. --accum-step sets the gradient accumulation steps, used to increase the effective batch size.
